# Development of a genetic toolset for the highly engineerable and metabolically versatile *Acinetobacter baylyi* ADP1

**DOI:** 10.1101/696302

**Authors:** Bradley W. Biggs, Stacy R. Bedore, Erika Arvay, Shu Huang, Harshith Subramanian, Emily A. McIntyre, Chantel V. Duscent-Maitland, Ellen L. Neidle, Keith E.J. Tyo

## Abstract

One primary objective of synthetic biology is to improve the sustainability of chemical manufacturing. Biological systems can utilize a variety of carbon sources, including waste streams that pose challenges to traditional chemical processing such as lignin biomass, providing opportunity for remediation and valorization of these materials. Success, however, depends on identifying microorganisms that are both metabolically versatile and engineerable. This has been a historic hindrance. Here, we leverage the facile genetics of the metabolically versatile bacterium *Acinetobacter baylyi* ADP1 to create easy and rapid molecular cloning workflows, a promoter library, ribosomal binding site (RBS) variants, and an unprecedented number of bacterial chromosomal protein expression sites and variants. Moreover, we demonstrate the utility of these tools by examining ADP1’s catabolic repression regulation, creating a strain with improved potential for lignin bioprocessing. Taken together, this work establishes ADP1 as an ideal host for a variety of sustainability and synthetic biology applications.

---

A critical element of sustainability is creating a closed and efficient carbon cycle by designing processes that utilize renewable resources and minimize or reclaim waste streams^1,2^. The emergence of synthetic biology has provided revolutionary new opportunities to perform sustainable and green chemistry^3^. In addition to operating at ambient reaction conditions and without the need for harsh reagents such as heavy metals, a key advantage for natural systems is their ability to utilize and adapt to a wide variety of feedstocks as carbon sources. This metabolic flexibility is exemplified by the conversion of waste C1 gases to ethanol with remarkable tolerance to real-time changes in feed gas content and quality using acetogens^4,5^. Such demonstrations highlight the potential for synthetic biology to expand and improve waste remediation processes.

To integrate additional waste streams, though, metabolically versatile microorganisms must be identified that are engineerable, capable of quickly undergoing many Design-Build-Test-Learn (DBTL) cycles. Recent advances in synthetic biology have identified numerous hosts with advantageous traits such as solvent tolerance^6^, novel metabolic capabilities^7^, and fast growth rates^8^, but progress with these hosts is often slowed by their genetic intractability. Unlike model organisms such as *Escherichia coli* and *Saccharomyces cerevisiae*, emerging hosts have comparatively few genetic tools and require lengthy DBTL cycles. Ideally, novel hosts would be identified that expand current functional capabilities *and* are easily manipulated.

For these reasons, the bacterium *Acinetobacter baylyi* ADP1 represents an exciting and ideal emerging synthetic biology host. ADP1 has a small genome, grows quickly (doubling time up to 35 minutes in rich medium)^9^, is naturally competent (i.e., capable of taking up DNA in standard growth conditions), and its native homologous recombination machinery enables facile allelic replacement^10,11^. Moreover, by way of the β-ketoadipate pathway, ADP1 is able to convert renewable, lignin derived aromatics to simple carbon building blocks^12,13^. This is significant as lignin is a major and notably underutilized component of non-food biomass^14^. Lignin’s heterogeneity and complexity have precluded simple approaches for its upgrading^15^, and metabolic engineering has been proposed as a solution^16^. Though other bacteria including *Pseudomonas* and *Rhodococcus* species have been identified with similar lignin consumption attributes to ADP1^12^, and advances have been made to their genetic toolsets^17–19^, these hosts can be challenging to engineer. ADP1 provides an opportunity to advance lignin bioprocessing through accelerated engineering cycles.

To validate this potential, we leveraged the natural competency and genetic tractability of ADP1 to establish simple and rapid cloning workflows that show notable reductions in experimental time compared to even established hosts such as *E. coli*. With these workflows, we created a promoter library, RBS variants, and characterized an unprecedented number of locations and promoter variants for chromosomal protein expression. To demonstrate the utility and reproducibility of these tools, we collaborated with an additional laboratory and examined the regulated consumption of aromatic monomers in ADP1. This work resulted in the creation of a strain with improved lignin bioprocessing attributes and shows that others can readily adopt these tools. Our work highlights ADP1’s amenability for biofoundry workflows^20^ and should establish ADP1 as a preferred option for lignin-based metabolic engineering, along with numerous other applications.

## RESULTS

### Evaluating and optimizing ADP1 transformations

To begin, we wanted to validate a simple standardized cloning workflow. Previous works established varied transformation protocols^9,10,21–23^, a complete single-gene deletion library^22^, and an *E. coli* compatible shuttle-vector (pBAV1k)^24^, but lacked consensus for molecular cloning steps. Owing to ADP1’s natural competency and native homologous recombination machinery, transformation workflows follow a general outline of: (1) incubation of ADP1 with transforming DNA in either liquid or on solid medium and (2) selection. Using a modified version of the available pBAV1k vector (**Supplemental Fig. 1**), we tested the simplest possible conditions (LB, liquid medium, short incubation times) and identified 30ºC, 3-hour incubation with as little as 25 ng of transforming DNA as highly efficient. Overnight incubations provided little to no improvement in transformation efficiency compared to 3 hours (**Supplemental Fig. 2**). In addition, incubations at 37ºC and shorter than 3 hours negatively impacted transformation rates (**Supplemental Fig. 3**). Using this workflow, we successfully transformed plasmid DNA, ligation products, Gibson products, and PCR products by natural competency (summarized in **Fig. 1**). Owing to ADP1’s growth rate (**Supplemental Fig. 4** and **5**), these protocols not only require much less experimental effort (∼10 minutes of experimental time) compared to *E. coli* transformation protocols but provide colonies on the same timeframe as *E. coli* (overnight).

**Figure 1.**
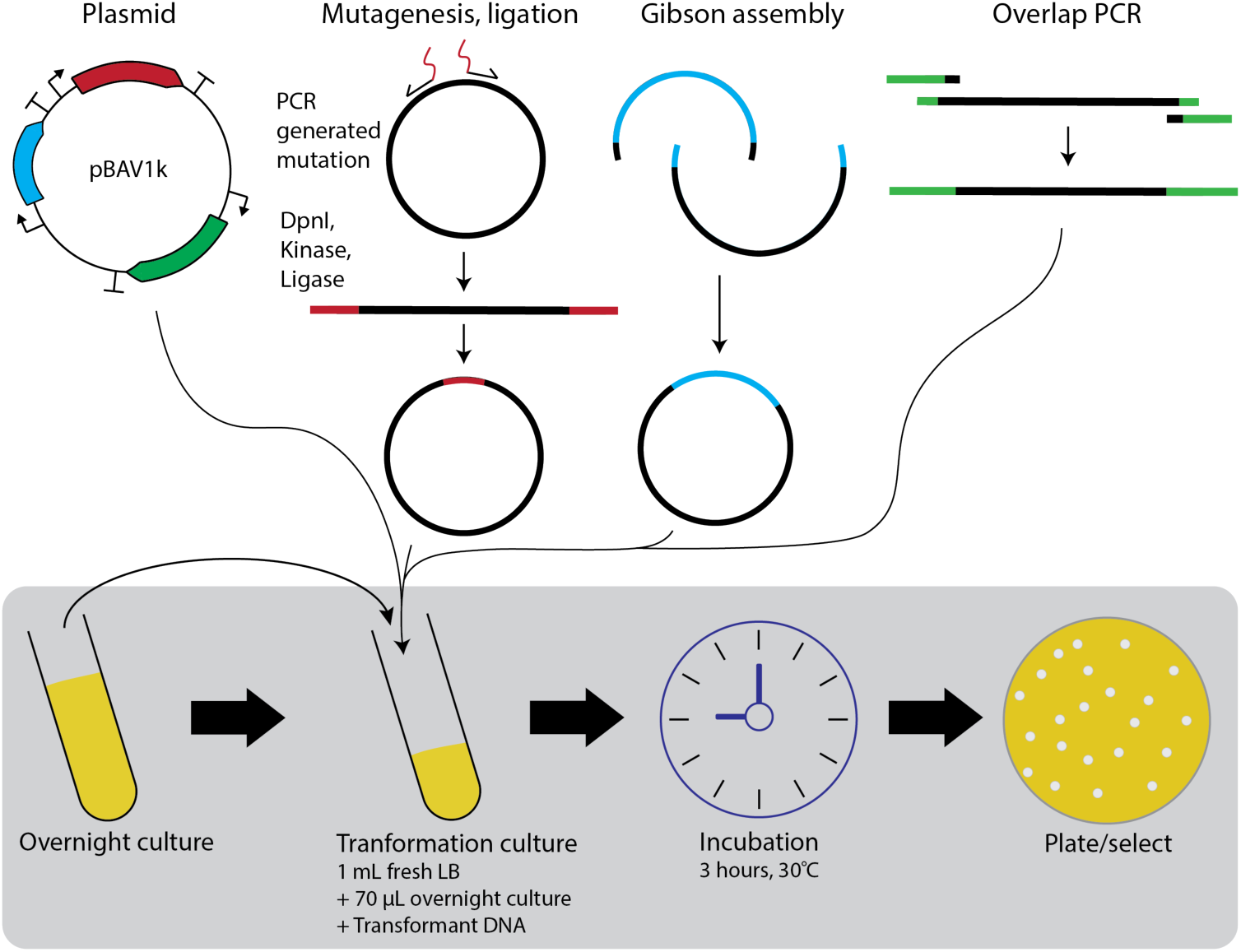
ADP1 cloning workflow. Transformation of ADP1 follows a simple process of sub-culturing an overnight culture grown in LB from a glycerol stock, adding transforming DNA, incubating for a short period of time (3 hours) at 30°C, and then selecting. With this workflow, we were able to transform plasmid DNA, ligation products, Gibson products, and linear PCR products.

**Figure 2.**
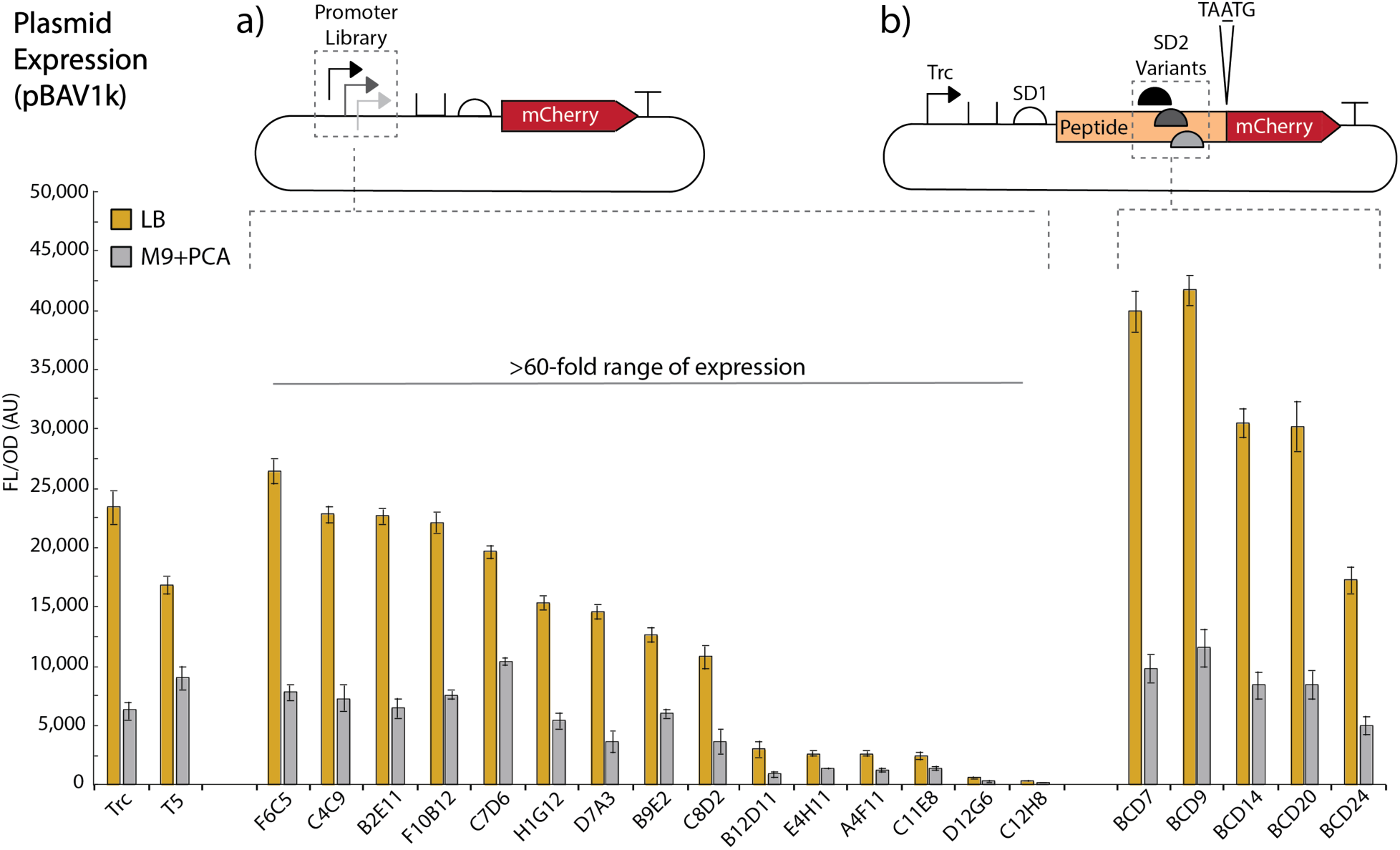
Plasmid expression of promoter library and RBS variants. Built by introducing “N” variability surrounding the −35 and −10 boxes of Trc in the context of pBWB162, our promoter library (a) shows 73-fold expression range for LB and 60-fold expression range for M9 with PCA. (b) Utilizing the BCD design to create RBS variants, we observe expression variance related to ribosomal binding strength. All experiments were performed as biological triplicate on three separate days (nine replicates), except LB expression of D12G6, which only had eight replicates. Error bars represent SEM.

**Figure 3.**
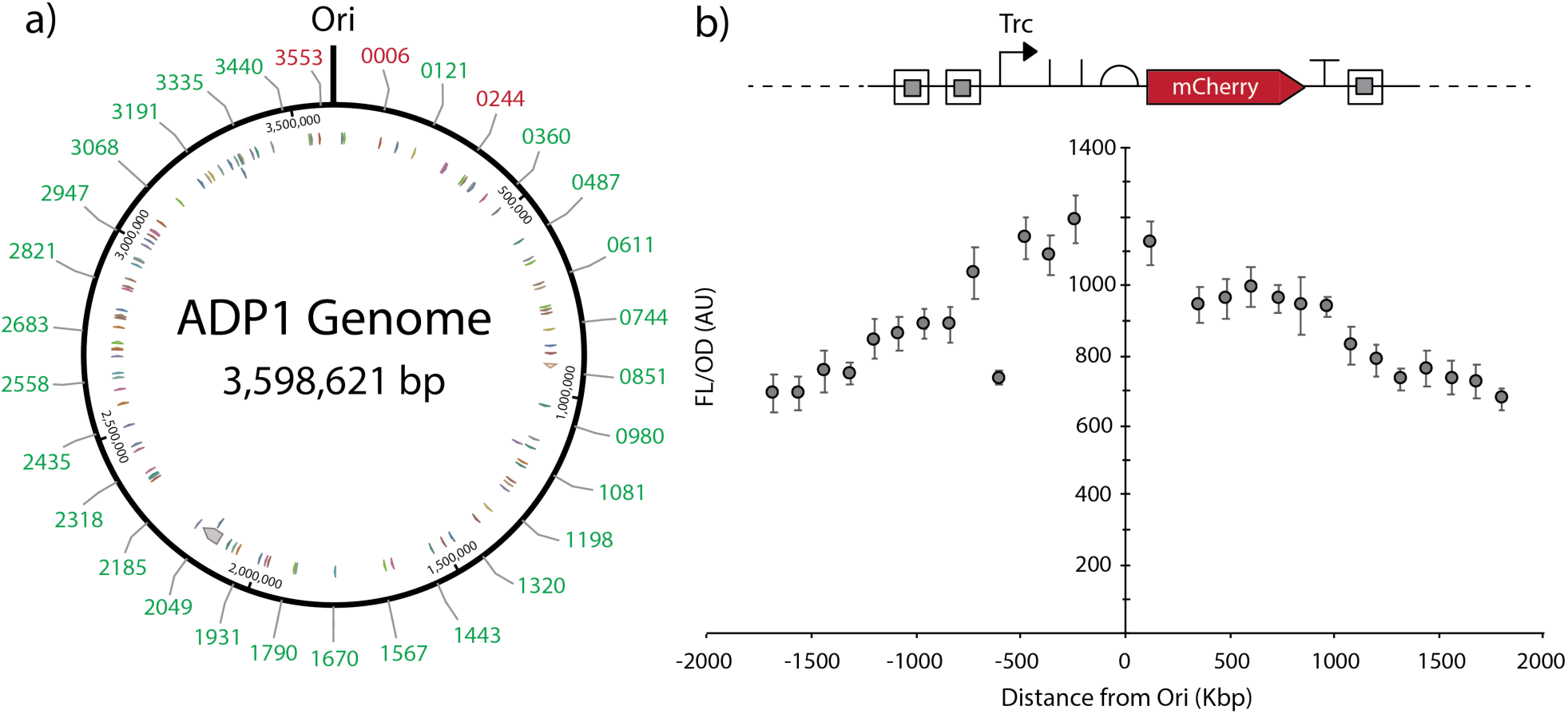
ADP1 chromosomal expression mapping. (**a**) Representation of the ADP1 genome, highlighting sites of integration by their ACIAD number. We integrated our insulated mCherry expressing cassette (pBWB206) (b) at 30 locations evenly spaced throughout ADP1’s genome, exactly replacing the gene in that location, unless its reading frame overlapped with an upstream or downstream gene. In those cases, the upstream or downstream gene was left intact. Expression orientation was always away from the origin of replication (Ori). Of the 30 integration sites, 27 (**a**, green numbers) produced a typical growth phenotype. These 27 sites were assayed for mCherry expression. (**b**) Expression shows an obvious correlation to distance from the Ori. One notable exception is ACIAD3068, which corresponds to *relA*, a stringent reponse related gene. While this integration did not impact growth, lack of stringent response regulation may impact protein expression. All cultures were carried out in M9 with 8 mM PCA and in biological triplicate on three separate days (nine replicates). Error bars represent SEM. Circular ADP1 genome representation originally generated using the NCBI ADP1 genome viewed by Benchling.

**Figure 4.**
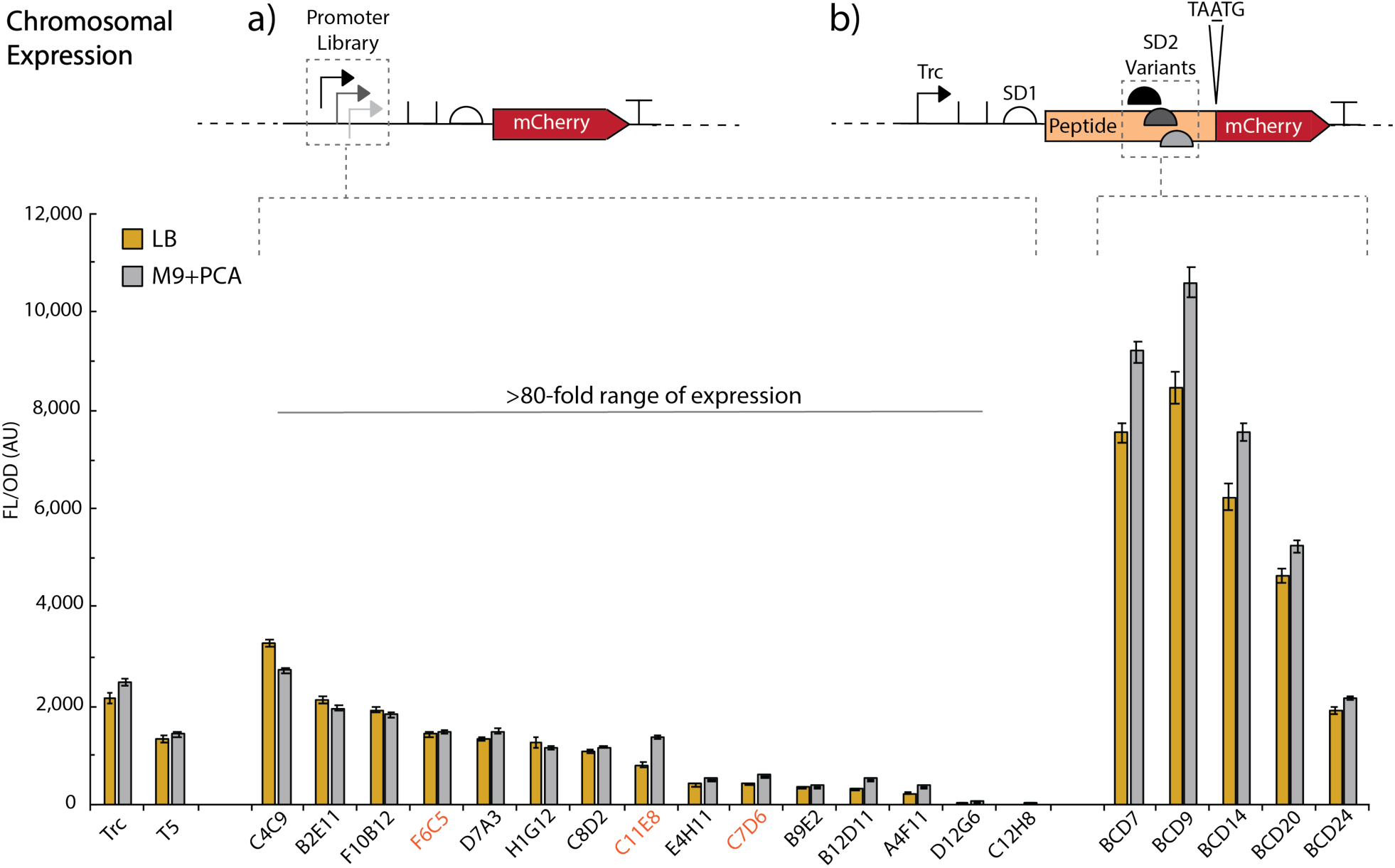
Chromosomal expression of promoter library and RBS variants. By integrating the entire library at ACIAD0980 (*vanAB*), we were able to examine the chromosomal expression behavior of our promoter library and set of BCD variants. (**a**) The promoter library performs similarly to plasmid expression, albeit with lower overall expression. However, the library still gives a >80-fold expression range for both LB and M9. In addition, M9 expression more closely matches LB expression on the chromosome compared to plasmid expression. (**b**) The BCD variants shows less of a decrease in expression when moved to the chromosome, particularly M9 BCD expression. A few promoter library variants behaved differently with respect to relative expression within the library, highlighted in orange. All experiments were run in biological triplicate on three separate days (nine replicates total). Error bars represent SEM.

Interestingly, when the shuttle vector used in this study was miniprepped from ADP1 there was ∼2-fold improvement in transformation efficiency compared to a miniprep from *E. coli* (**Supplemental Fig. 6**), with approximately 41,000 cfu/µg for this 5 kb plasmid using our workflow. We hypothesize that methylation patterns may help ADP1 recognize DNA as “self.” In addition, we found that ADP1 provides strong plasmid yields from a standard miniprep kit (>3-4 µg for a 5 mL overnight culture) and, as suggested in a previous work^25^, that WT ADP1 harbors no native plasmids (**Supplemental Table 1**). Worth noting, to obtain clarified lysates for a standard miniprep kit, we needed to extend the centrifugation step after neutralization to two >10 minutes spins at 17,000 *g* (max speed) to obtain suitable DNA purity. In all, ADP1 transformation requires no specialized preparation or purchasing of competent cells, involves a single 10-minutes hands on step compared to five steps of equal or greater complexity for *E. coli* heat shock, gives high transformation efficiencies, does not extend time for colony growth, and could be easily parallelized with simple liquid handling equipment.

### Creating a promoter library

Precise control of gene expression is critical for engineering microorganisms, so we next moved to construct expression tools^26^. We first sought to validate previous work^27^ with a set of common synthetic biology promoters. In addition, we wanted to compare these promoters to the bacterial components of a newly developed set of broad-spectrum promoters^28^, along with potentially strong-expressing native ADP1 promoters chosen based on ADP1 transcriptomic data^29^. Each promoter was put into the context of pBWB162, a derivative of pBAV1k with an mCherry reporter, which was first tested for potential readthrough and inducibility and found to have minimal leakiness and to be stably induced with 1 mM IPTG (**Supplemental Figs. 7** and **8**).

Testing this group of promoters (**Supplemental Fig. 9**), we found that bacterial consensus-based promoters (Trc, Tac, T5) all gave strong expression. This is perhaps expected as ADP1 carries an RpoD/σ70 homolog and previous *in vitro* work showed that *E. coli* σ70 RNA polymerase gave transcription of an ADP1 gene with its native promoter and transcription factor^30^. T7 did not give expression, in contrast to a previous study^27^. The synthetic broad-spectrum promoters gave expression, but the relative expression strength of the four promoters did not match order of strength found in the original work^28^ (**Supplemental Fig. 9**). Though the native ADP1 promoter for *gltI*, potentially the strongest native promoter based on transcriptomic data^29^, gave strong expression, it was still not as strong as Trc (**Supplemental Fig. 9**). Because of this, in combination with Trc’s promoter architecture being well understood while *gltI’s* is not, we chose to construct a promoter library based on Trc. Leveraging ADP1’s natural competency, promoter library construction followed a simple workflow (**Supplemental Fig. 10**). Mutagenic primers were used to introduce variability flanking and in between the bacterial consensus −35 and −10 boxes, and clones with a range of expression strength were isolated (**Supplemental Fig. 11**, full description of library construction in **Supplemental Methods**). Sequences for the promoter library can be found in **Supplemental Table 2** and full plasmid sequence files are available with the **Supplemental Materials**.

We sought to identify genetic parts with robustness to different conditions, and as promoter expression characteristics can change depending on the growth medium, we rescreened the promoters in both LB and unmodified M9 with 8 mM protocatechuate (PCA) as the sole carbon source (**Fig. 2**). Unless otherwise stated, all M9 cultures used PCA as the only carbon source. We chose PCA because it is a common intermediate in the breakdown of lignin via the β-ketoadipate pathway. Our 15-member promoter library shows 73-fold expression range in LB and 60-fold expression range in M9 (**Fig. 2a**).

Next, to test the effect of RBS modulation on expression, we picked a range of RBS variants from the previously developed bicistronic design (BCD) library^31^. Briefly, this design involves two RBSs, where the first RBS initiates the translation of a leader peptide for which the final nucleotide of the stop codon overlaps with the start codon of the gene of interest (TA<u>A</u>TG). This design is proposed to overcome RBS performance variability due to mRNA secondary structure by utilizing the ribosome’s helicase activity. These RBS/BCD variants were successfully applied in ADP1 and showed a 2.4-fold range of expression in LB and 2.3-fold range of expression in M9 (**Fig. 2b**). Interestingly, certain promoter and BCD variants performed differently in LB compared to M9-based expression, highlighting the need to screen in each medium before applying genetic parts to metabolic engineering contexts. Examples include Trc and T5 switching order of strength, BCD7 and BCD9 switching order of strength, and C7D6 and B9E2 showing relatively greater expression strength in M9. Overall, our promoter library and RBS variants together provide access to >100-fold range of expression strength in ADP1.

### Mapping genomic integrations

For industrial metabolic engineering applications, genomic integration of a metabolic pathway is typically desirable for stable long-term expression without a selective pressure^32^. Because ADP1 readily incorporates DNA into the genome, this was straightforward. Accordingly, we tested a variety of chromosomal locations in ADP1 for their capacity for heterologous protein expression. To demonstrate the ease of ADP1 genomic integration, we chose 30 non-essential gene sites evenly spaced throughout the genome, based upon a previous complete single-gene knockout study^22^. In this approach our knock in cassette directly replaced the non-essential gene (GenBank files with the reporter cassette in a given loci are available with the **Supplemental Materials**). This scale of genomic integration is greater than any other bacterial chromosomal expression screen, including previously constructed *E. coli* and *Bacillus subtilis* libraries^33,34^.

To generate homologous recombination cassettes, we utilized an overlap PCR strategy to incorporate 500 bp of homology on either side of an insertion cassette (**Supplemental Fig. 12**). Multiple attempts to use either 50 or 100 bps of flanking homology, introduced by PCR, for knock ins using this 3.7 kb insertion cassette gave no transformants in our hands. In addition, the λ-red system^35^ appeared too toxic in ADP1, as multiple different plasmid constructs for gam-β-exo expression could not be stably maintained. To prevent transcriptional readthrough from biasing analyzed expression, our integration cassette bore three additional terminators to act as insulators (pBWB206). These insulators consisted of two upstream terminators and one downstream terminator following the vector’s pre-existing terminator. Insulators were chosen based on a previously designed orthogonal terminator library^36^. To some capacity the insulators functioned as attenuators, as their inclusion lowered plasmid-based expression 20-fold compared to the parent plasmid pBWB162 (**Supplemental Fig. 13**). After overlap PCR assembly, linear PCR product was either directly added to the medium or gel extracted and then added to the medium, following the established transformation workflow.

Of the 30 sites tested, 27 had correct transformants from a single attempt, identified by screening eight colonies by colony PCR. We tested these sites for expression, plotting expression with respect to distance from the origin of replication (**Fig. 3** for M9, **Supplemental Fig. 14** for LB). For consistency, the orientation of all expression cassettes was away from the origin of replication, and interestingly the (-) strand generally gave higher expression (**Fig. 3b**). This was also true for two locations where we integrated the expression cassette in both orientations (**Supplemental Fig. 15**). Expression was found to have a clear inverse correlation to the distance from origin of replication. Overall, this mapping provides numerous and diverse integration sites for stable and effective protein expression in ADP1.

### Integration of promoter library and RBS variants into the genome

To further highlight the ease of genomic manipulation in ADP1, we integrated the entire promoter library and set of BCD variants at a single locus (ACIAD0980, genes *vanAB*). These knock ins were constructed with a similar overlap PCR strategy as the genomic integration mapping cassettes, but on a cassette without additional insulators. Upon screening, the promoter library was found to give >80-fold range of expression for both LB and M9 (**Fig. 4a**). M9-based expression was closer to LB expression when tested on the genome compared to the plasmid. In fact, the BCD variants actually showed greater expression per OD_600_ in M9 than LB (**Fig. 4b**). There was a more significant decrease in expression by moving to the chromosome for the promoter library than for the BCD variants (**Supplemental Figs. 16-19**). This is perhaps due to the overall lower copy numbers of mRNA being generated from chromosomal expression and the ability of the strong BCDs to maintain high translation per mRNA in both circumstances. Worth noting, three promoters in the library changed their position in terms of expression strength (F6C5, C11E8, C7D6, all highlighted with orange on **Fig. 4a**) upon chromosomal integration. To our knowledge, this is the most thorough comparison between plasmid and chromosomal integration expression in the literature.

Finally, to ensure we could perform successive genomic integrations, we tested a counter-selection workflow. We found we needed to provide strong *sacB* expression (pBWB290, **Supplemental Fig. 20**) and to perform sucrose counter-selection at room temperature to achieve adequate counterselection pressure. Upon screening 50 counter-selected colonies by colony PCR, we found 8% of the 50 colonies screened contained the desired vector at the locus of interest and were clonally pure after restreaking. In addition, we found this vector capable of marker-less heterologous protein expression (**Supplemental Fig. 21**).

### Tool application for lignin valorization

To demonstrate the applicability of these tools and their reliability and reproducibility in different labs (at Northwestern University and at the University of Georgia), we studied aspects of ADP1’s catabolite repression. The ultimate goal is to enable simultaneous consumption of lignin-derived aromatic compounds in mixtures of pre-treated biomass^12,16^. This type of consolidated bioprocessing depends on engineering bacteria for rapid production of valuable end-products and may require overriding catabolite repression, which imposes preferential carbon source utilization^37,38^. In ADP1, all aromatic compounds are consumed via one of two branches of the β-ketoadipate pathway (**Supplemental Fig. 22**). ADP1 preferentially consumes benzoate, degraded via the catechol branch of this pathway before *p*-hydroxybenzoate (POB) degraded via the protocatechuate (PCA) branch^39^. Studies of benzoate and POB co-metabolism can, therefore, serve as a proxy for more complex mixtures, including lignin hydrolysates.

BenM and CatM are transcriptional regulators that respond to a catechol-derived metabolite, muconate, to activate *ben* and *cat* genes^40^. Although BenM and CatM may repress *pca* genes, their exact role in the benzoate-mediated delay of POB catabolism remains unclear^39^. One hypothesis is that repression of a *pca* gene (*pcaK*, which encodes a POB transporter), prevents the formation of PCA by lowering POB uptake (**Supplemental Fig. 23**). The new gene expression tools were used to investigate this hypothesis. Specifically, we tested how the following affect co-consumption: (a) constitutive expression of PcaK, (b) ablation of BenM/CatM binding sites that could repress *pcaK* transcription, or (c) deletion of *benM* and *catM* with constitutive expression of genes for benzoate degradation to allow benzoate and POB to be degraded in the absence of both regulators (**Supplemental Fig. 24**).

To facilitate uptake of POB in the presence of benzoate, the PcaK transporter was constitutively expressed under the control of several synthetic promoters such that its transcription would not be repressed by BenM or CatM. Despite using different strength promoters for PcaK expression, these attempts to increase POB transport did not accelerate its degradation (**Supplemental Fig. 25**). Sequential consumption of benzoate before POB was observed as with the wild type (**Fig. 5bii** compared to **5bi)**. Second, to alleviate BenM and CatM-based repression of *pcaK*, the repressive binding sites upstream of the *pca* operon were scrambled. Again, this change was not sufficient to allow co-consumption (**Fig. 5biii** and **Supplemental Fig. 26**). However, deletion of BenM and CatM, coupled with constitutive expression of benzoate degrading enzymes, led to co-consumption of POB and benzoate (**Fig. 5biv**). Collectively, these results imply BenM and CatM act to repress POB degradation by multiple methods, and only their removal alleviates this type of catabolite repression. These studies, and additional results and discussions in a **Supplemental Note**, demonstrate the utility and ease of implementation for these tools, including the chromosomal use of multiple synthetic promoters in the same strain. This methodology not only contributes to improved understanding of multiple carbon source consumption but also delivers a strain that may prove useful in lignin valorization.

**Figure 5.**
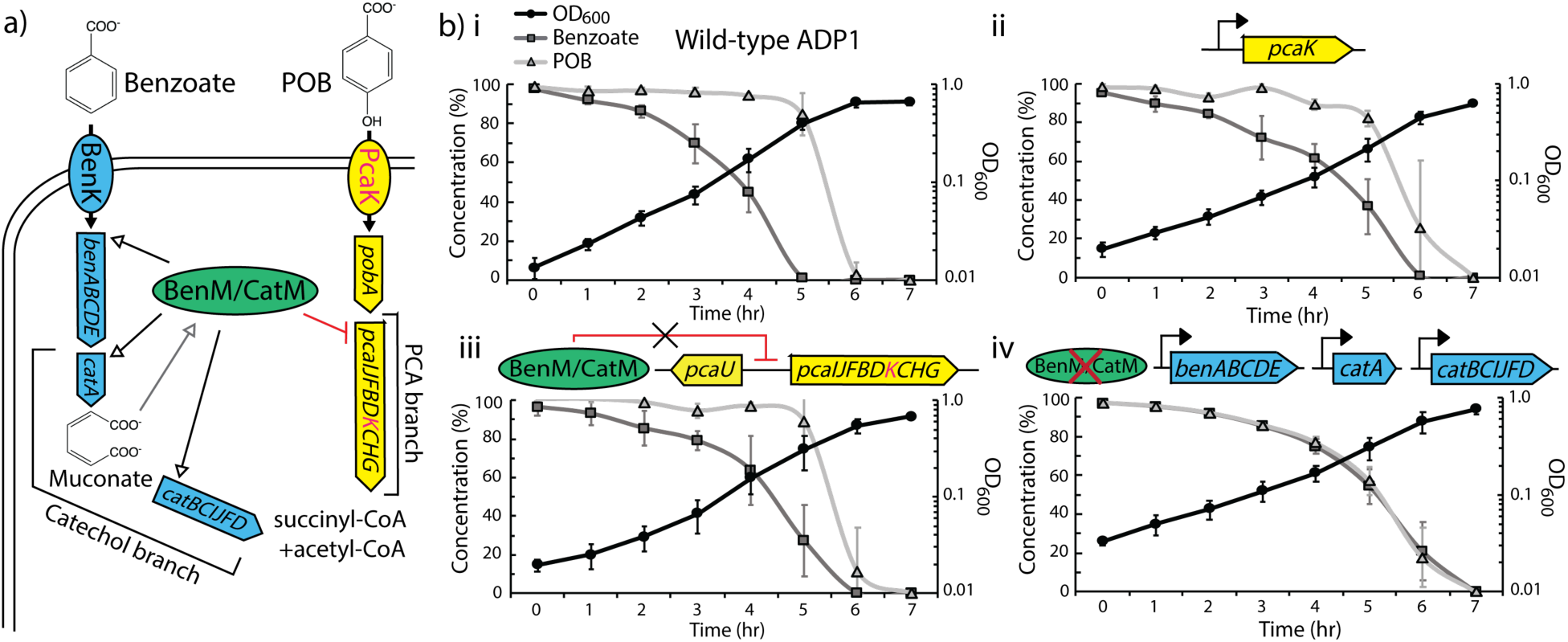
Lignin consumption engineering. (**a**) Schematic depicting regulation and metabolism of *p*-hydroxybenzoate (POB) and benzoate, where the two molecules enter the PCA and catechol branches of the β-ketoadipate, respectively, to eventually be funneled to central carbon metabolism. After being imported across the cell membrane, POB is degraded by *pob* and *pca* genes, but the *pca* genes can be repressed by BenM/CatM. Whereas, benzoate is consumed by *ben* and *cat* genes, which are activated by BenM/CatM. (**b**) Schematic showing the consumption patterns of different strains. First, **i)** shows wild-type ADP1 consumption where benzoate is consumed prior to POB. **ii)** Shows consumption where the transporter *pcaK* is placed under constitutive expression with a synthetic promoter. This consumption remains sequential. **iii)** Shows consumption pattern under the condition where the binding site for BenM/CatM upstream of the *pca* operon has been scrambled. This consumption still occurs sequentially. Finally, **iv)** shows an engineered strain where *benM* has been inactivated and *catM* has been deleted, and where the catechol branch enzymes, which are normally activated by BenM and CatM, are under constitutive expression with synthetic promoters (T5 for *catA*, F6C5 for *catBCIJFD*). This strain shows simultaneous co-consumption. Experiments run as five biological replicates. Error bars represent standard deviation.

## DISCUSSION

The cumulative tools developed in this study, along with other recent works that have established an additional plasmid vector^41^, CRISPR elements^41^, and ADP1 chromosomal evolution tools^42^, effectively equip ADP1 as a model organism for lignin-based metabolic engineering and other synthetic biology applications. ADP1 has significant advantages over other microorganisms for DBTL cycles. The simplicity associated with natural transformation and facile allelic replacement lends itself to automation, liquid handling, and incorporation into biofoundry workflows^20^, especially as advances in DNA synthesis have alleviated the bottleneck of gene synthesis. Modifying the genome (as oppose to transforming with plasmids) is straightforward, highlighted by the >50 chromosomal integrations carried out in this study. As the chromosomal tools developed in this study even exceed far more established hosts such as *E. coli* and *B. subtilis*, we expect this work to facilitate ADP1’s use as a host for stable and robust engineering.

Our validation in two different laboratories indicates the tool set is reproducible and scientifically rigorous. Further highlighting the importance of these tools is the ease with which regulatory hypotheses were tested, revealing new information about preferential carbon source utilization and generating strains for use in lignin valorization. Milestone strains from this study have been deposited for easy access on Addgene, and cloning files are available with the **Supplemental Materials**. In addition, we have attempted to document approaches that were unsuccessful as a service to others that might work with ADP1.

Because of its ease of engineering, the metabolic versatility of ADP1 may expand to metabolic engineering applications beyond lignin-derived aromatics to more traditional glucose-based processes. As ADP1 exclusively utilizes the cofactor-generating Entner-Doudoroff pathway while maintaining high growth rates, as opposed to *E. coli’s* preferred use for maximal growth of the Embden–Meyerhof–Parnas glycolytic pathway^43^, there may be benefit to using ADP1 to produce products with heavy cofactor needs. Indeed, Santala and coworkers have already provided demonstration of glucose-based metabolic engineering in ADP1^44^. Ultimately, we envision this host will be adopted beyond the lignin community and will play a role in the future of sustainability and synthetic biology.

## METHODS

Methods and any associated references are available in the online version of the paper.

## ACKNOWLEDGMENTS

We would like to thank Dr. Gregg T. Beckham, Dr. Isabel Pardo, and Dr. Christopher W. Johnson for their helpful conversations about this manuscript. We gratefully acknowledge the contributions of Melissa Tumen-Velasquez to the construction and design of some ACN strains and pBAC plasmids. This work was supported by NIH Biotechnology Training Grant (T32-GM008449-23 to B.W.B., E.A.), the National Science Foundation (collaborative grant MCB 1614953 to K.E.J.T. and MCB 1615365 to E.L.N.), and the Northwestern University NUSeq Core Facility.

## AUTHOR CONTRIBUTIONS

B.W.B, E.L.N., and K.E.J.T. conceived and designed the study. B.W.B., S.H., and S.B. performed cloning. E.A. and H.S. performed growth curve experiments. B.W.B. performed fluorescent expression experiments. S.R.B., E.A.M., and C.V.D.-M. performed co-consumption experiments. B.W.B., S.R.B., E.L.N., and K.E.J.T. wrote and prepared the manuscript.

