## Supplemental Materials for "Development of a genetic toolset for the highly engineerable and metabolically versatile *Acinetobacter baylyi* ADP1"

- Supplemental Table 3: Plasmid for aromatic compound catabolism studies........ 34-36
- Supplemental Table 4: ADP1-derived strains for aromatic compound catabolism studies........ 36-37

Supplemental Note........ 38-40
Supplemental Files Description........ 41
Supplemental References........ 42-43

### Methods

***Cell culturing***. In this study wild-type *Acinetobacter baylyi* (ADP1)[1,2] and *Escherichia coli* DH5α were used for all cloning and cell culture work. *E. coli* K12 MG1655 was used for growth comparison tests. ADP1 was used for all expression tests and genetic part validation. Culture tube cultivations were carried out 30ºC, 250 rpm, unless otherwise noted. Plate-based cultivations were carried out at 30ºC, 150 rpm. Cultivations were run in Fisher's LB Broth Miller (LB) or Difco M9 Minimal Medium (M9). M9 was supplemented with various carbon sources as indicated. Kanamycin (kan) at 25 µg/mL was used for antibiotic selection. Agar plates were made with 15 g of Teknova Agar per liter of medium. Agar plates needed to be well dried before ADP1 plating to get well-defined circular colonies. For growth tests, optical density was measured using a Shimadzu UV-1800 UV-Vis Spectrophotometer at an absorbance of 600 nm ($OD_{600}$). Glycerol stocks were stored at -80ºC and were created by adding 750 µL of overnight cell culture medium to 250 µL of 60% (v/v) glycerol for a final glycerol concentration of 15% (v/v). Growth tests were carried out in LB, M9 with 0.4% glucose, and M9 with 5 mM each *p*-coumarate and ferulate. Cultures of 25 mL were inoculated at an $OD_{600}$ of 0.05 and grown at 30ºC or 37ºC and 250 rpm in 250 mL unbaffled shake flasks. Cultures were grown in biological triplicate, except for in the case of ADP1 growth in M9 minimal medium with 0.4% glucose (w/v), which was carried out in 9 replicates.

***Cloning***. PCRs were carried out with PrimeSTAR Max 2x master mix from Takara Bio, following manufacturer instructions for annealing temperatures and extension times using the three-step protocol (denature, anneal, extend). An Excel file with a list of all primers used in this study can be found with the supplementary information [**Supplemental File 2 (Primers)**]. In addition, cloning files representing the final parts used for this study can found with the supplemental information. General cloning, primer design, and *in silico* Gibson assembly design were carried out with Benchling. Gibson assemblies were run using NEB 2x Gibson master mix, following manufacturer protocols. All mutagenic primers were designed on the NEBaseChanger web interface (https://nebasechanger.neb.com/). All blunt end ligations following primer-based mutagenesis were carried out using on gel purified DNA and the "KLD" mix from the NEB Q5 Site-Directed Mutagenesis Kit, following manufacturer protocols.

PCR templates for colony PCRs were obtained by picking individual colonies and resuspending them into 20 µL of nuclease-free water, while patching them to a separate agar plate for secondary validation and to obtain clonally pure strains. For colony PCR, master mixes were prepared and ~10 µL colony PCRs were run with 9.8 µL of the master mix and 0.5 µL of the water in which the colony had been resuspended as the template. All final genomic integrations were sequenced via their colony PCR product.

***Fluorescent measurement experiments***. Fluorescent measurements were taken with a Synergy H1 Microplate Reader from BioTek. mCherry readings were taken at both the top and bottom with excitation at 585 nM and emission at 615 nM. Gain was set at 100. The read height was set at 7 mm. Reported fluorescent values were exclusively taken from the top measurements. Plate reader cell density measurements were taken by reading the $OD_{600}$. Plate reader experiments were carried out in biological triplicate, except for the pBAV1k promoter library, pBAV1k RBS (BCD) variants, chromosomal integration mapping, chromosomally integrated promoter library, and chromosomally integrated RBS (BCD) variants, which were carried out as biological triplicates run on three separate days (nine replicates total).

To prepare plate reader fluorescent experiments, overnight cultures were carried out in sterile 0.4 mL 96-well flat bottom plates (Fisher Cat. No. 267576) at 30ºC and 150 rpm, scraping glycerol stocks for inoculation into 200 µL of LB with appropriate antibiotic. Sub-culturing (1:100 dilution) was carried out the following morning by successive dilutions of first 40 µL of LB overnight culture added to 160 µL of the final medium (LB or M9, with corresponding antibiotic and carbon source, which was 8 mM protocatechuate for the M9 medium) to make an initial 40:200 (1:5) dilution plate. Then, 10 µL of this 1:5 dilution was added to 190 µL of the final medium (1:100 final dilution). The final cultures were run in sterile 0.4 mL optical bottom black 96-well plates. After 3 hours, these cultures were induced with 1 mM IPTG (5 µL of 40 mM IPTG stock). Fluorescent measurements were taken the next morning after gentle resuspension by pipetting up and down, being careful to not introduce any bubbles to the culture. All cultures were covered with Sterile AeraSeal films. Films were replaced with a fresh sterile film each time they were removed (for induction and for fluorescent measurements).

Fluorescence per $OD_{600}$ [FL/OD (AU)] was calculated by first subtracting the values of fluorescence and $OD_{600}$ for the blank medium from all wells. Next fluorescence was divided by $OD_{600}$. Following, the wild type ADP1 average FL/OD value for that medium context was subtracted from all wells. Averages and standard deviations or standard error were calculated, as indicated.

***Natural transformation***. Natural transformation was carried out by first inoculating wild-type ADP1 from a glycerol stock into a 3 mL overnight culture, grown at 30ºC and 250 rpm in a culture tube. The following day, 70 µL of this culture was added to 1 mL of fresh LB medium in a culture tube and incubated for 3 hours along with the transforming DNA (plasmid, PCR product, Gibson product, ligation product). Transforming DNA did not need to be cleaned up by gel extraction or PCR clean up kit before direct addition to the ADP1 culture, and as little as 25 ng of transforming DNA added to the 1 mL of culture was sufficient to obtain transformants. After the 3 hours, 150 µL of the medium was plated on the selective plate (either kanamycin or sucrose), spread with glass beads, and allowed to dry briefly before moving to the incubator.

***Promoter library creation***. The promoter library was created using the above natural transformation workflow. A single set of mutagenic primers was purchased with “NNN” diversity 5 bp upstream of the bacterial consensus -35 box (TTGACA), in the 17 bp in between the -35 and -10 box (TATAAT), and for the 5 bp after the -10 box [**Supplemental File 2 (Primers)**, primers BWB361, BWB362]. A single PCR was run with the pBWB162 vector as the template. The product was gel extracted and added into the KLD NEB mix (DpnI, kinase, ligase) and allowed to react for 5 minutes at room temperature. This product was then directly added into the ADP1 culture for transformation. After 3 hours, 150 µL of culture was plated on two separate plates. Plates were incubated at 30ºC. The next day, 96 colonies were picked and inoculated into a 96 well plate for fluorescence screening.

***Genomic integration***. For the genomic integration mapping, an insulated integration cassette was first constructed by the addition of two upstream terminators and one additional terminator downstream. These insulators were added by successive PCR mutagenesis via overhangs to create pBWB206.

Once the genomic integration cassette was created, knock in cassettes were constructed by overlap PCR. Overlap PCRs were run in two stages. First, three PCRs were run to obtain two ~500 bp oligos with homology matching the up and downstream regions of the ADP1 gene to be knocked out, respectively, and the insulated genomic integration cassette (**Supplemental Fig. 12**). Each part had 20-30 bp of overhang homology to the part to which it would be assembled by overlap PCR. Approximately 1 µL of each gel extracted part was added to a 45 µL PCR reaction and run for 14 cycles to create the full-length knock in cassette. Following, outer primers were added to amplify the full-length knock in cassette for an additional 19 cycles. Knock in cassettes were either directly added to the ADP1 medium or gel extracted first. Approximately 5 µL of either PCR product, gel extracted or not, was added to the medium. The same workflow was used for the promoter library and RBS/BCD variant integrations, but these cassettes did not bear the insulator regions.

***Counter-selection***. Counter-selection was conducted using the newly created pBWB290 vector as the initial integration cassette. Full length knock in cassettes were generated by overlap PCR as described above for the genomic integrations. After initial integration, colonies were patched both for kanamycin resistance and sucrose sensitivity. Only colonies that showed kanamycin resistance and sucrose sensitivity were carried forward for colony PCR validation. Counter-selection was carried out with 12.1% sucrose, NaCl-free LB agar plates. To create sucrose plates, LB components were first autoclaved and sterile filtered sucrose was added after autoclaving. After plating or patching, counter-selection was carried out at room temperature. Hits from patching were screened by colony PCR.

For counter-selection, a 3 mL ADP1 culture was incubated overnight in LB with kanamycin. The next morning, this culture was spun down (4,000 rpm, 10 minutes, 4ºC), the medium was decanted, and then the pellet was resuspended in 3 mL of fresh LB without antibiotic. 70 µL of this resuspended culture was added to 1 mL of LB along with ~5 µL of the marker-less integration cassette. This culture was incubated for 3 hours, and then 150 µL was plated on the 12.1% sucrose LB agar plates and left at room temperature overnight. In general, a full 24 hours of room temperature growth was necessary to see colonies large enough to be picked. (Note: If the initial sacB/kan integration was not first patched on sucrose to identify integrants with sucrose sensitivity, counter-selection would show a hazy lawn with colony growth on top of this lawn. These colonies growing on the lawns were typically false positives.) However, if sucrose sensitive variants were first identified and validated by colony PCR, it was possible to avoid a lawn "background" on the sucrose plate. For sucrose plates with clear colonies, these colonies were patched on sucrose and kanamycin LB agar plates. Colonies that grew on sucrose and had kanamycin sensitivity were then screened by colony PCR. Colony PCR hits were then sent for sequencing.

***Media and growth conditions for studies of aromatic compound consumption.*** Because catabolic studies were designed for comparison with past results, the media and growth conditions matched previous investigations. A defined minimal medium was used for culturing *A. baylyi* strains[3]. This medium was supplemented with 2 mM benzoate and 2 mM POB for carbon source consumption experiments and 20 mM pyruvate for natural transformation culturing. *E. coli* XLI Blue cells (Stratagene) were used as plasmid hosts and grown in LB[4]. All cultures were incubated at 37°C with aeration (shaking at 250 rpm).

***Strain and plasmid construction for studies of aromatic compound consumption.*** DNA for plasmid construction was PCR-amplified with a high-fidelity polymerase, PrimeSTAR Max (Takara Biosciences) or Phusion (New England Biosciences), using the primers listed in **Supplemental File 2 (Primers)**. Plasmids were assembled using an overlapping sequence method conducted *in vivo* in *E. coli* XL1-Blue competent cells (Stratagene)[5]. In some cases, standard restriction digestion was followed by ligation (Quick Ligation Kit, New England Biosciences). Plasmids are described further in **Supplemental Table 3**. Chromosomal changes in *A. baylyi* were constructed by allelic replacement[6, 7] and are listed in **Supplemental Table 4**. After transforming recipients with donor DNA, desired transformants were selected for drug resistance or growth in the presence of sucrose, as previously described[7]. Alternatively, cells were screened to detect transformants that lost the ability to metabolize POB by plating on solid minimal medium with 2 mM succinate and 2 mM POB. Cells that can grow on both carbon sources form larger colonies than those that can grow on succinate but not POB. The small colonies were then patched to test for growth on 5 mM POB as the sole carbon source. Isolates that were unable to grow on POB as the sole carbon source were further evaluated. All plasmids and strains were confirmed by PCR and/or regional DNA sequencing.

To create strains with altered PcaK expression, the *pcaK* coding sequence was first deleted from its normal operonic position (to generate ACN2451). ACN2451 was then used as the transformation recipient for linearized plasmids carrying *pcaK* controlled by different constitutive promoters. Each *pcaK* allele was introduced between convergently oriented chromosomal genes (*quiA* and ACIAD1717) with independent transcription ensured by a transcriptional terminator engineered downstream of *quiA*, as depicted in **Supplemental Fig. 25**. A drug resistance cassette (ΩK*52468*) downstream of the plasmid-borne *pcaK* was used to select transformants that had undergone allelic replacement.

***Creating random changes in the CatM/BenM binding site.*** To prevent CatM and BenM from binding to a site upstream of *pcaU* (**Supplemental Fig. 23**), changes were introduced by PCR with a mutagenic primer (SRB368) designed to eliminate two conserved features of this site important for interactions with CatM and BenM:[8] 1) a conserved sequence (T-$N_{11}$-A), and 2) a small region of dyad symmetry surrounding this sequence. The mutagenic primer was synthesized with random nucleotides (NNNN) in place of the chromosomal sequences in ADP1 at positions 1,708,958-1,708,961 and 1,708,969-1,708,972 (NCBI entry NC_005966). PCR products were cloned to generate plasmids (pBAC1794, pBAC1795, pBAC1797, pBAC1798, pBAC1799, and pBAC1800) in which there is an ΩK cassette downstream of *pcaU.* These plasmids were linearized and used to transform ADP1 to create strains in which the plasmid-borne alleles replaced the corresponding wild-type regions (ACN2529, ACN2496, ACN2498, ACN2530, ACN2526, and ACN2499, respectively). These strains were initially selected by the drug resistance conferred by ΩK. DNA sequencing confirmed the specific mutations that were introduced in these plasmids and strains. In all cases, the two key features of the wild-type BenM/CatM binding site (noted earlier) were eliminated (see **Supplemental Tables 3** and **4**). These strains retained the ability to use POB as the sole carbon source.

***Alteration of sequences to prevent homologous recombination between two different synthetic promoters used on the genome of a single strain.*** As depicted in **Supplemental Fig. 24**, DNA is identical in the regions surrounding the critical promoter sequences of different transcriptional constructs in the tool kit. To use two promoters (F6C5 and T5) to control the transcription of two adjacent *cat*-gene chromosomal regions, the DNA surrounding the promoter sequence of one promoter was altered to avoid the possibility of recombination between identical sequences. The *lac* promoter from pBTL-2[9] was changed by site directed mutagenesis with primers SRB441 and SRB442 to generate the F6C5 promoter on plasmid pBAC1682.

***Carbon source consumption experiments.*** For each strain tested for aromatic compound catabolism, cells were first grown in 5 mL minimal medium in test tubes overnight and sub-cultured 100 µl into 5 mL fresh medium to be grown overnight again. Then 1 mL of the second culture was used to inoculate 50 mL fresh medium in a 250 mL Erlenmeyer flask from which hourly samples were taken. For experiments displayed in **Figure 5**, the inoculating cultures were grown on 2 mM POB. For experiments displayed in **Supplemental Figs. 25** and **26**, the inoculating cultures were grown on 2 mM benzoate and 2 mM POB. After experimental samples were taken, cells were removed by filtration, and the supernatant was analyzed to monitor metabolites by high performance liquid chromatography (HPLC) as previously described[10, 11]. Retention times for POB and benzoate were 6 min and 13.8 min respectively. Data were plotted with the 0 time-point defined as being one hour before depletion of any carbon source was first detected. Experiments were performed in biological triplicate. The optical density was measured at a wavelength of 600 nm on a spectrophotometer (Beckman DU 640 or Eppendorf Biophotometer Plus 6132).

#### Supplemental Figures

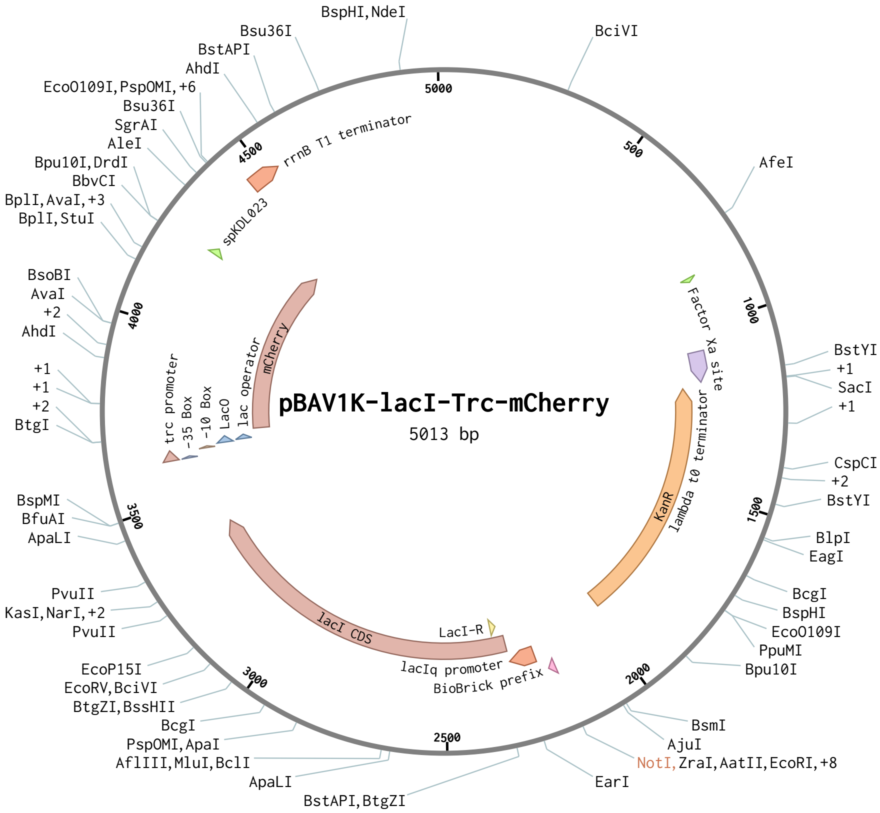

**Supplemental Figure 1.** *Plasmid map for pBWB162*. Map shows the modified version of pBAV1k, with lacI-Trc-mCherry, used as the basis for all fluorescent expression tests in this study. (Figure prepared with Benchling.)

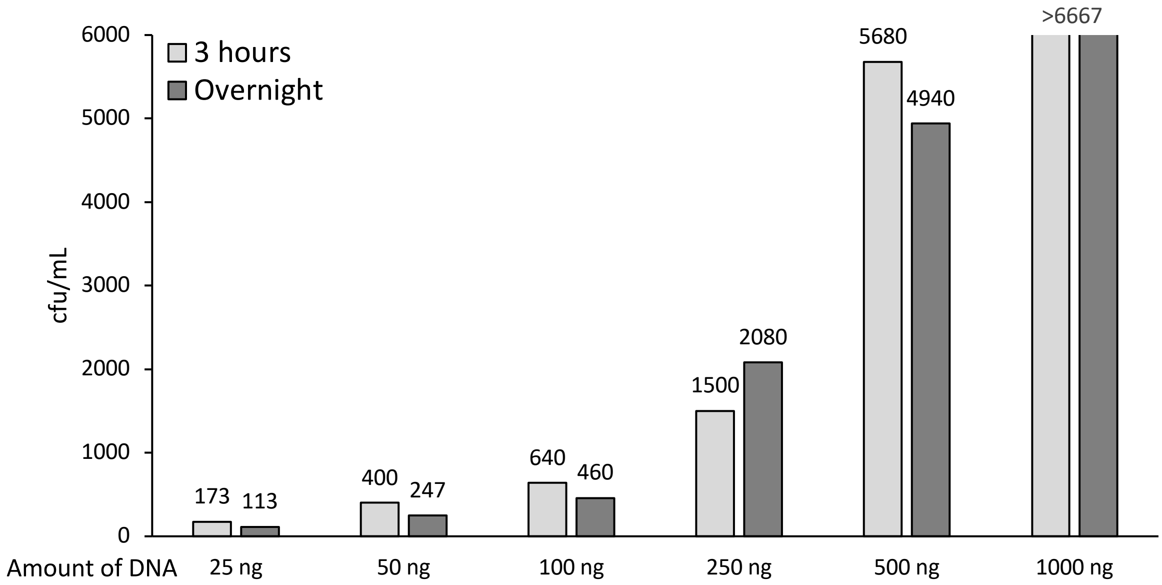

**Supplemental Figure 2.** *pBAV1k ADP1 transformation DNA concentration and timing comparison*. Figure shows colony forming units per mL of LB medium for a range of DNA concentrations used for pBAV1k plasmid transformation. As can be seen, useful transformation is achieved with as little as 25 ng of plasmid DNA added to the 1 mL of fresh medium inoculated with 70 μL of overnight culture previously inoculated by glycerol stock.

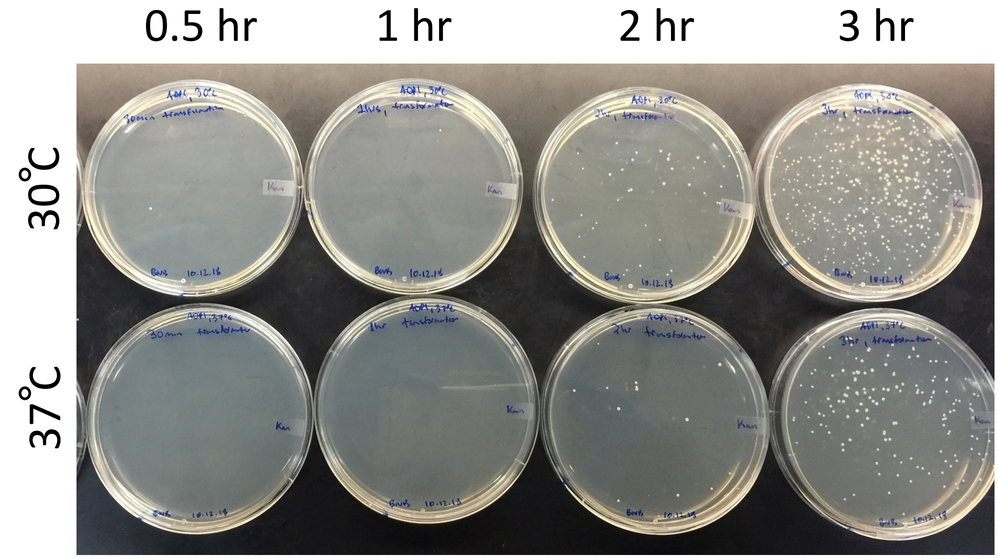

**Supplemental Figure 3.** *Comparing temperature and timing of ADP1 pBAV1k transformation.* Using 75 ng of pBAV1k to transform ADP1, inoculating 70 µL of overnight culture into 1 mL of fresh LB, cells were grown at both 30ºC and 37ºC and plated at 0.5, 1, 2, and 3 hours to compare transformation efficiency. As can be seen, times shorter than 3 hours result in a significant decrease in transformation efficiency. In addition, growing cells at 37ºC is detrimental to transformation efficiency for all times.

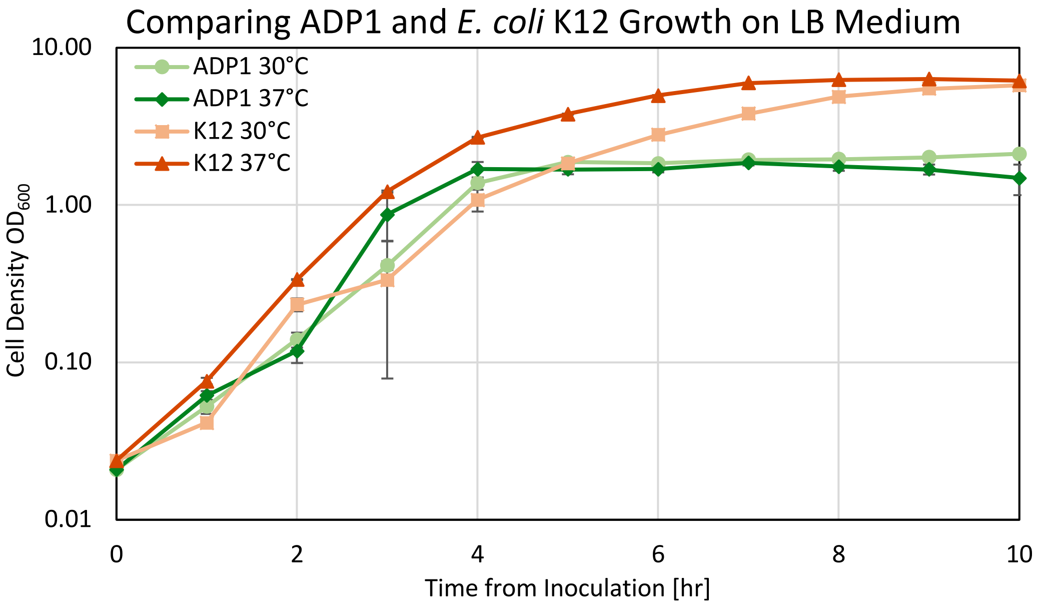

**Supplemental Figure 4**. *Comparison of ADP1 and E. coli K12 growth in LB at 30ºC and 37ºC*. As can be seen, during the initial growth stages on LB, ADP1 has a comparable growth curve to *E. coli* K12 before entering stationary phase at around 5 hours. ADP1's growth rate at 30°C is 1.09 $hr^{-1}$ with a doubling time of 38 min. ADP1 growth at 37ºC is 1.1928 $hr^{-1}$ with a doubling time of 35 min. *E. coli* K13 growth rate at 30ºC is 0.972 $hr^{-1}$ with a doubling time of 43 min. *E. coli* K12 growth rate at 37ºC is 1.3293 $hr^{-1}$ with a doubling time of 31 min. Error bars are for standard deviation of biological replicates.

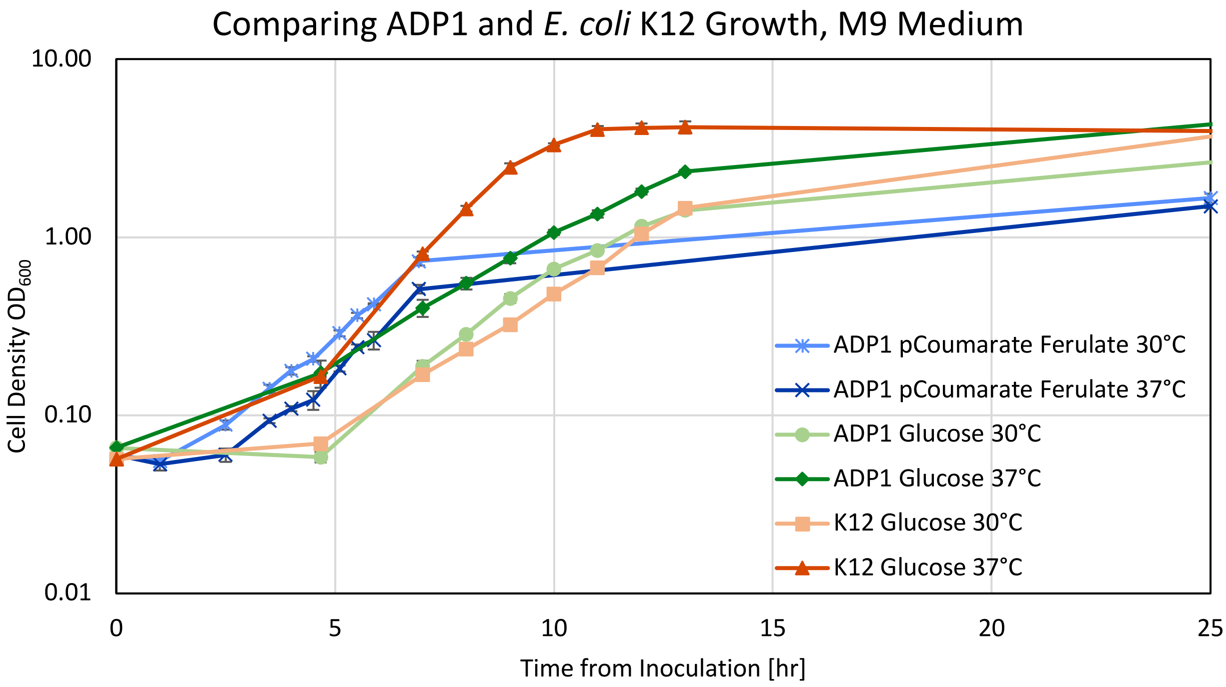

**Supplemental Figure 5.** *Comparison of ADP1 and E. coli K12 growth in minimal medium (M9) at 30ºC and 37ºC.* As can be seen, though *E. coli* K12 reaches stationary phase at 37ºC more quickly, ADP1 is able to achieve a higher overall OD. *p*-coumarate and ferulate were both provided at an initial concentration of 2 mM. Glucose was provided at 0.4% w/v. ADP1 growth on *p*-coumerate and ferulate at 30ºC was 0.5177 $hr^{-1}$ with a doubling time of 80 min, and at 37ºC the growth rate was 0.5791 $hr^{-1}$ with a doubling time of 72 min. ADP1 growth rate on glucose in minimal medium at 30ºC was 0.427 $hr^{-1}$ with a doubling time of 98 min, and for 37ºC the growth rate was 0.364 $hr^{-1}$ with a doubling time of 115 min. The growth rate for *E. coli* K12 in minimal medium on glucose at 30ºC was 0.291 $hr^{-1}$ with a doubling time of 152 min, and for 37ºC the growth rate was 0.591 $hr^{-1}$ with a doubling time of 71 min. Error bars are for biological replicates.

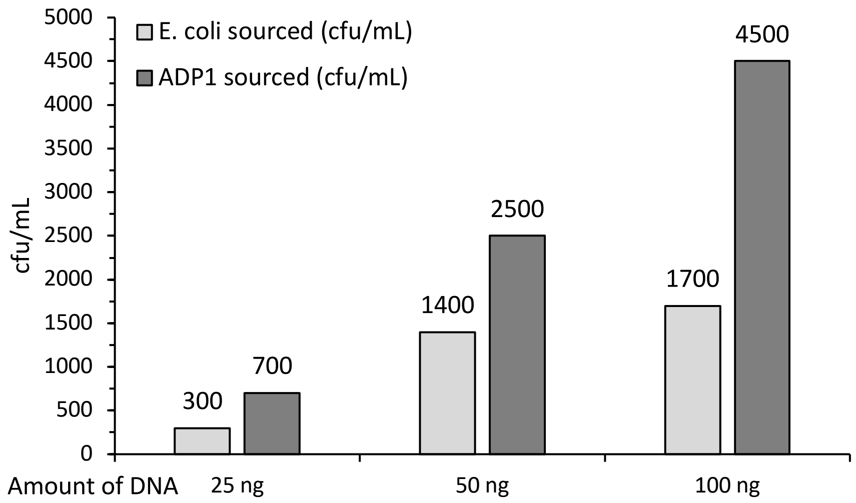

**Supplemental Figure 6.** *ADP1 transformation from E. coli and ADP1 prepped pBAV1k*. ADP1 was transformed with varying amounts of either *E. coli* or ADP1 miniprepped pBAV1k plasmid. As can be seen, ADP1 prepped DNA has a higher efficiency of transformation (greater number of colony forming units) for each concentration, giving an average of approximately 2.25x greater transformation efficiency.

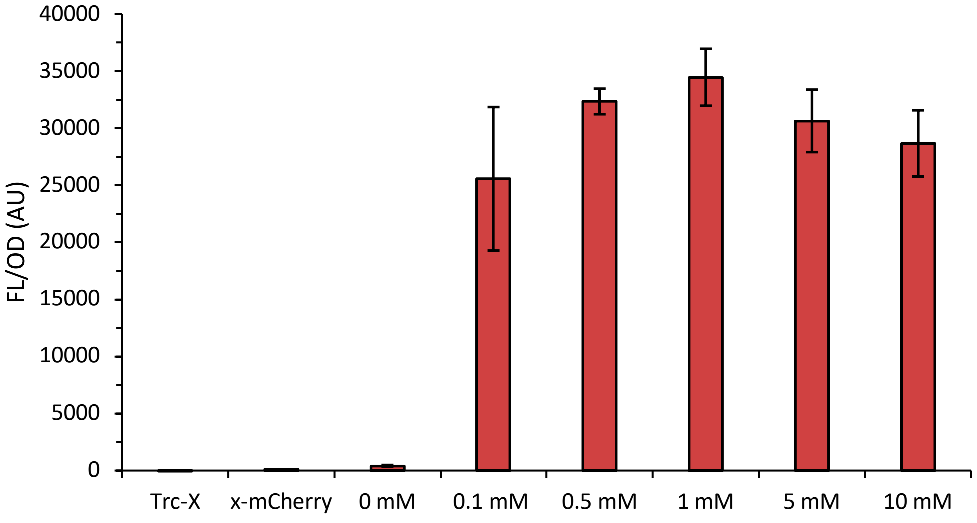

**Supplemental Figure 7.** *Readthrough and inducibility tests of pBWB162.* Using the newly constructed pBWB162 and two controls lacking either the mCherry reporter or promoter, inducibility and readthrough were tested. As can be seen, this vector has minimal readthrough or "leakiness," demonstrating the observed fluorescence is due to the promoter and true induction. Titrating inducer shows stable induction above 0.5 mM IPTG, and greatest induction at 1 mM IPTG. All experiments were run in biological triplicate in 96 well plates in LB. Error bars are standard deviation.

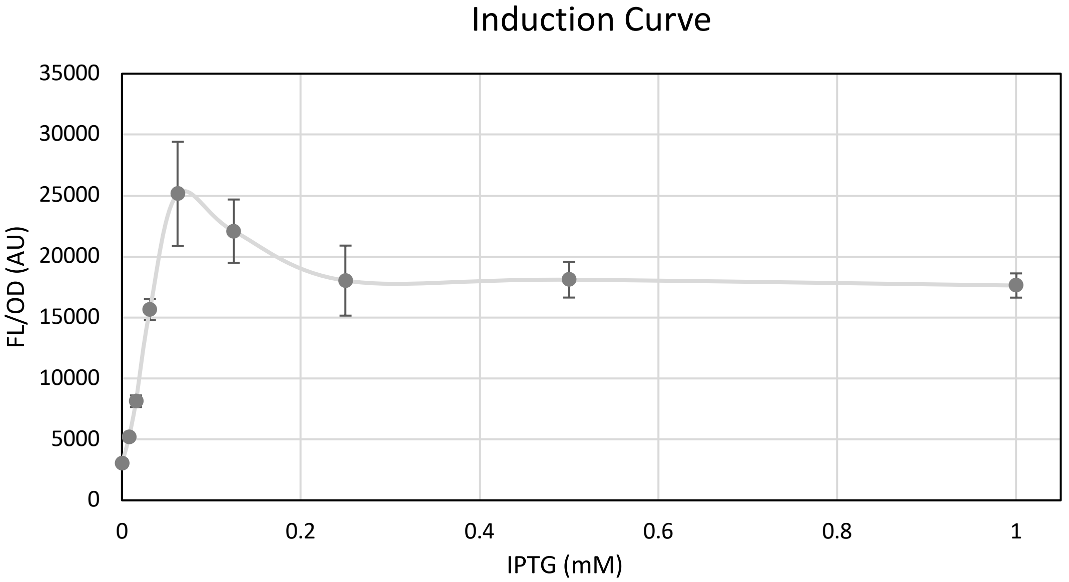

**Supplemental Figure 8.** *pBWB162 IPTG induction curve*. Again, consistent (low error bar) induction doesn't occur until 0.5 mM IPTG and the lowest error bars for stable high-expression induction are at 1 mM ITPG. Experiments run in biological triplicate in 96-well plates in LB. Error bars are standard deviation.

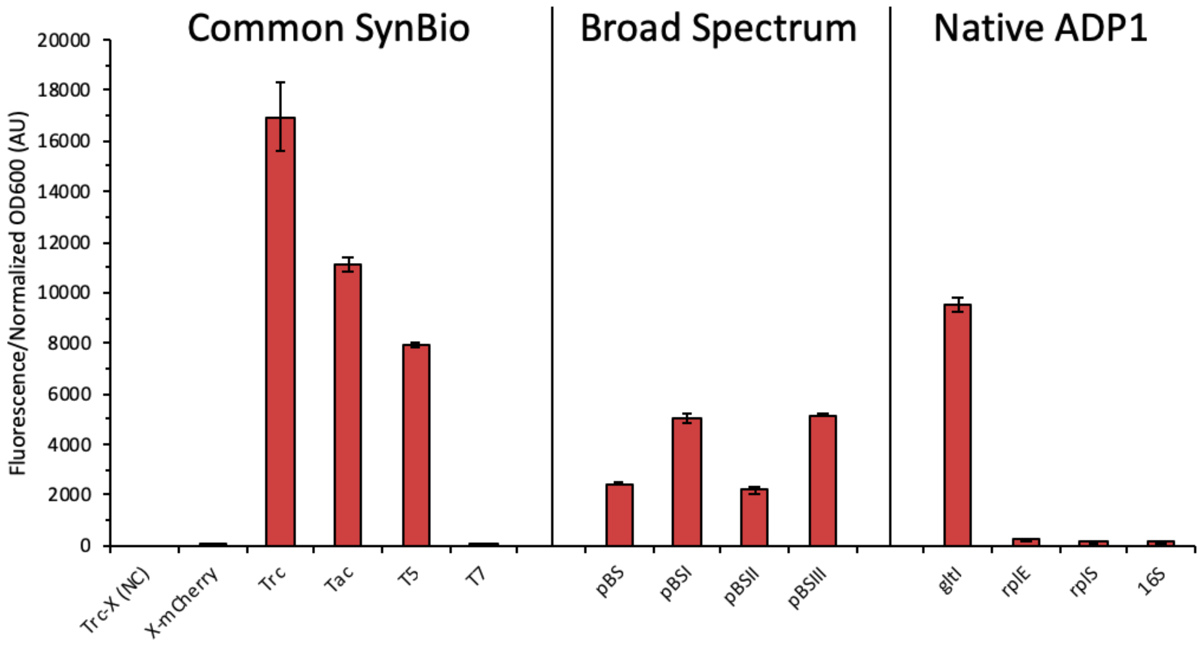

**Supplemental Figure 9.** *Initial promoter screening*. An initial group of promoters was screened in ADP1 to examine performance. Those based on the bacterial consensus promoter sequence (Trc, Tac, T5) show strong expression, as expected owing to the similarity of the *rpoD*/σ70 homologs in ADP1 and *E. coli*. Similarly, the bacterial portion of a group of broad-spectrum promoters from a recent work[12] show expression, however not in the order and strength expected. Finally, using the 300-500 upstream bases 5' to genes with known high mRNA abundance in ADP1,[13] the gene with the highest transcriptomic expression from previous work (*gltI*) gives the strongest expression on par with T5 and below Trc. Other native promoters did not show strong expression. All were run as biological triplicates in 96-well plates in LB. Error bars are standard deviation.

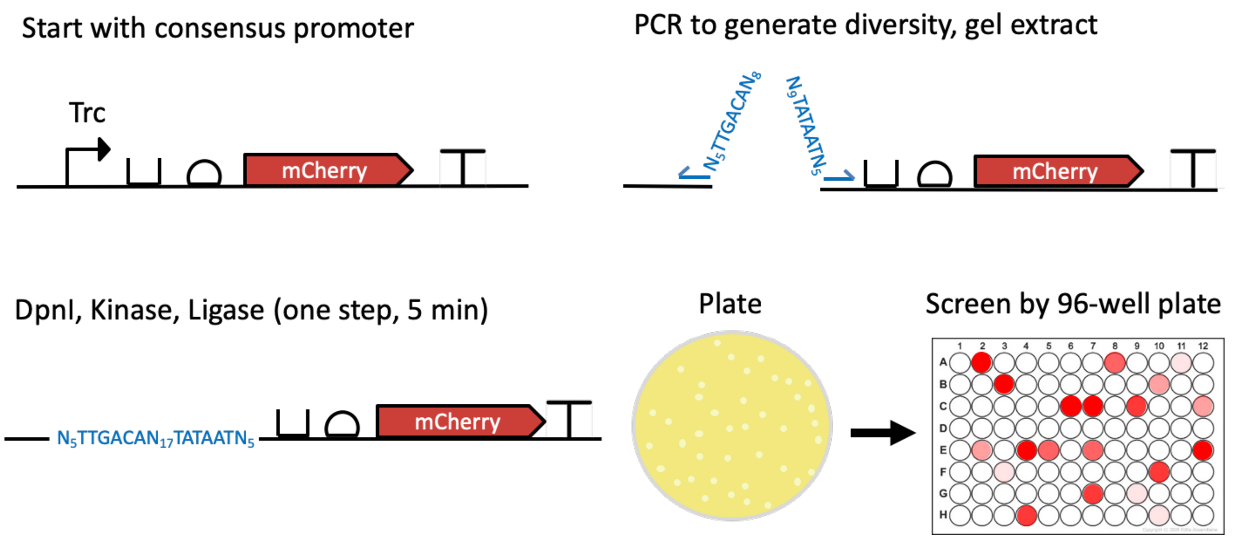

**Supplemental Figure 10.** *Workflow for promoter library creation*. Workflow shows the primer-based mutagenesis workflow for the creation of the promoter library. Built from pBWB162, the bacterial consensus -35 and -10 boxes were left intact, while the 5 bp upstream, 17 bp intervening, and 5 bp downstream were allowed to vary with any "N." Transformants were plated, picked, and assayed in 96-well plate format. Variants with expression signal above noise were carried forward for future experiments.

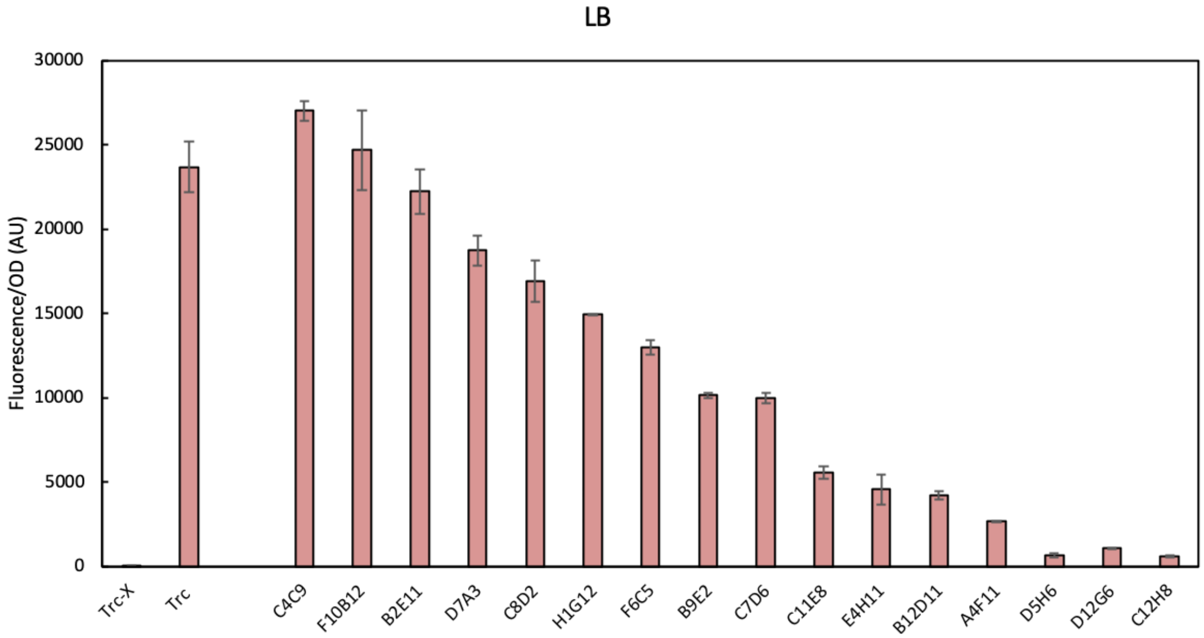

**Supplemental Figure 11.** *Initial screening of promoter library built from pBWB162 mutagenesis in LB*. After transformation in ADP1, colonies were picked and screened for mCherry expression in 96-well plate format (as depicted in Supplemental Fig. 10). Those with stable expression were archive and then re-screened to give this subset of 16. Of these 16, D5H6 was removed from further testing as it was prone to mutation, with the vector becoming mutated twice. D5H6 is shown in this iteration of the library, as it was used in the co-consumption work, and this data gives reference to its expression strength as a promoter. The sequence for D5H6 is included in Supplemental Table 2. Cultivations were run in biological triplicate. Error bars represent standard deviation of biogical triplicate.

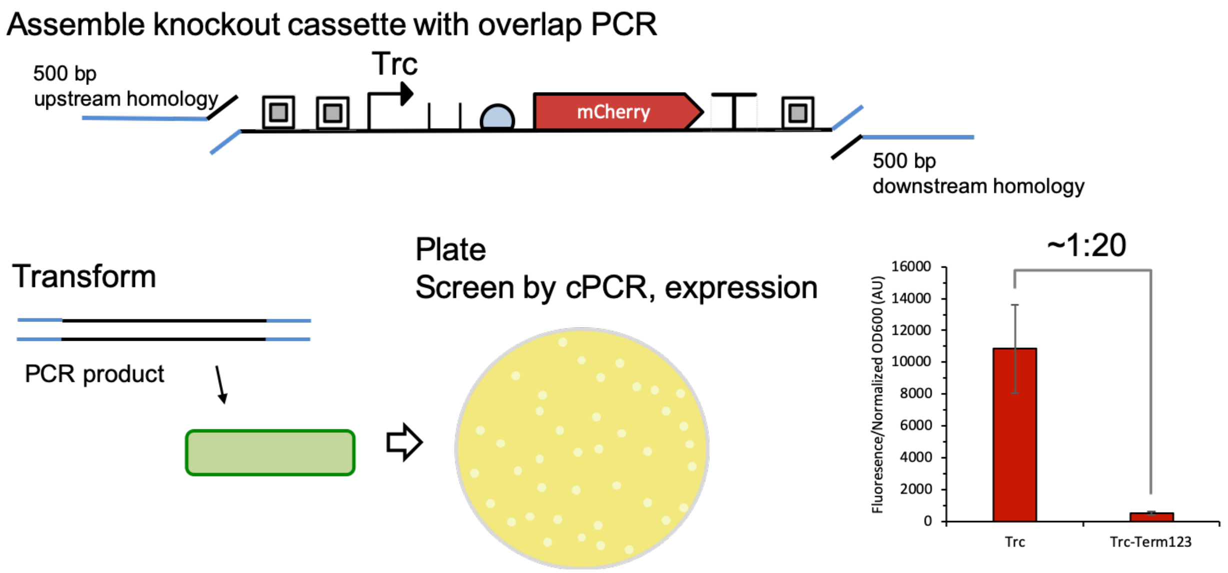

**Supplemental Figure 12.** *Workflow for overlap PCR-based knockouts*. Knockouts were accomplished using overlap PCR products carrying approximately 500 bps of homology flanking both ends of the knock in cassette. PCR was carried out in two-stages, first the flanking ~500 bp homology regions were PCR amplified off the ADP1 genome with 20-50 bp of homology to the knock in cassette. Similarly, the knock in cassettes, which contained flanking insulators, were PCR amplified off the template plasmid with 20-50 bp of overlap with the homology region. Overlap PCR was then carried out in two stages, first 14 cycles with just the three overlap PCR parts, then 19 cycles with the addition of outer primers capable of amplifying the full-length cassette. This overlap PCR product was then either gel extracted or directly added into the ADP1 natural transformation workflow.

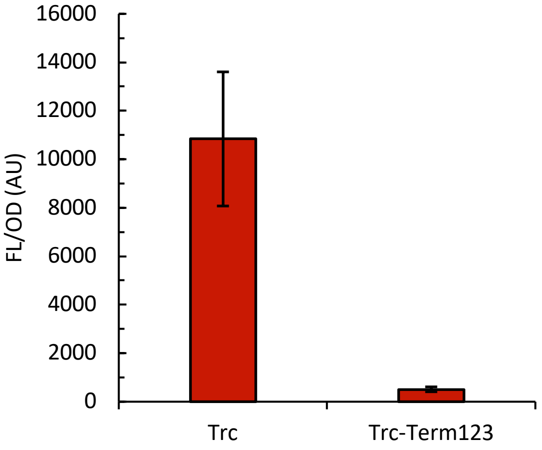

**Supplementary Figure 13.** *Expression of insulated pBWB206*. As can be seen, plasmid based expression of the insulated vector is greatly reduced (approxmiately 1/20$^{th}$ of the expression) compared to its predecessor with no insulators. In this graph Trc is analogous to pBWB162 and Trc-Term123 is analogous to pBWB206. Experiment was performed in biological triplicate and the error bars represent standard deviation.

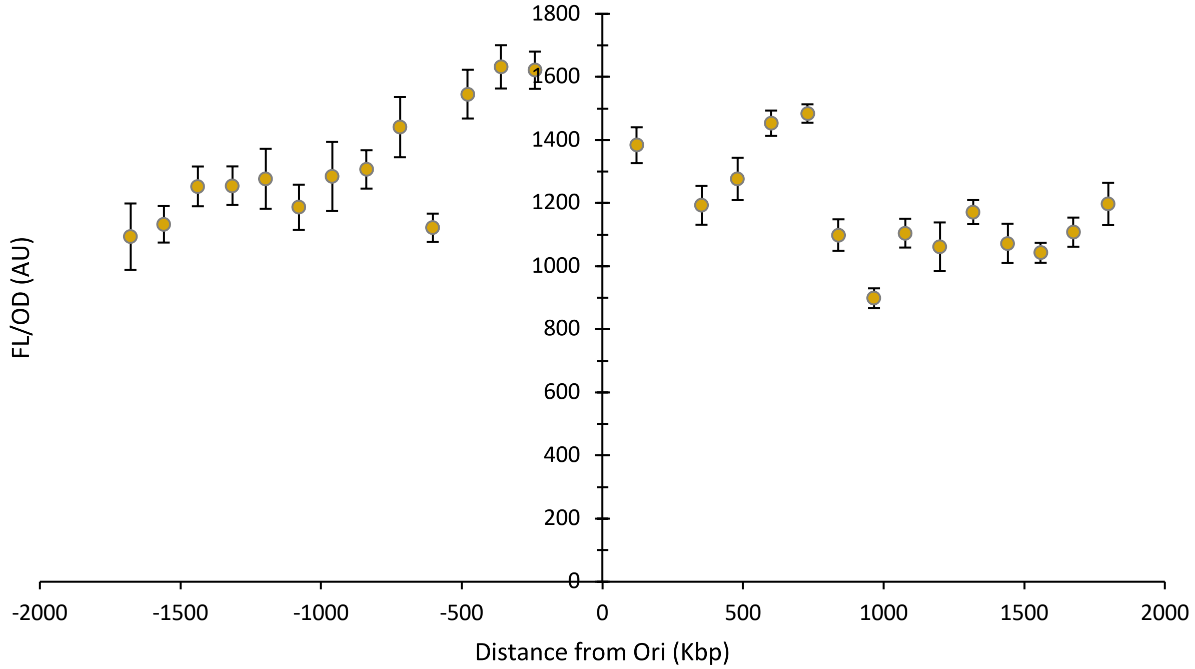

**Supplemental Figure 14.** *Chromosomal mapping expression in LB*. Similar to the M9 screen, there is a weak correlation of expression to distance from origin of replication. Secondly, the – strand (left side) seems to give higher expression on average. All experiments were performed in biological triplicate on three separate days to give (nine total replicates). Errors bars are for standard error of measurement.

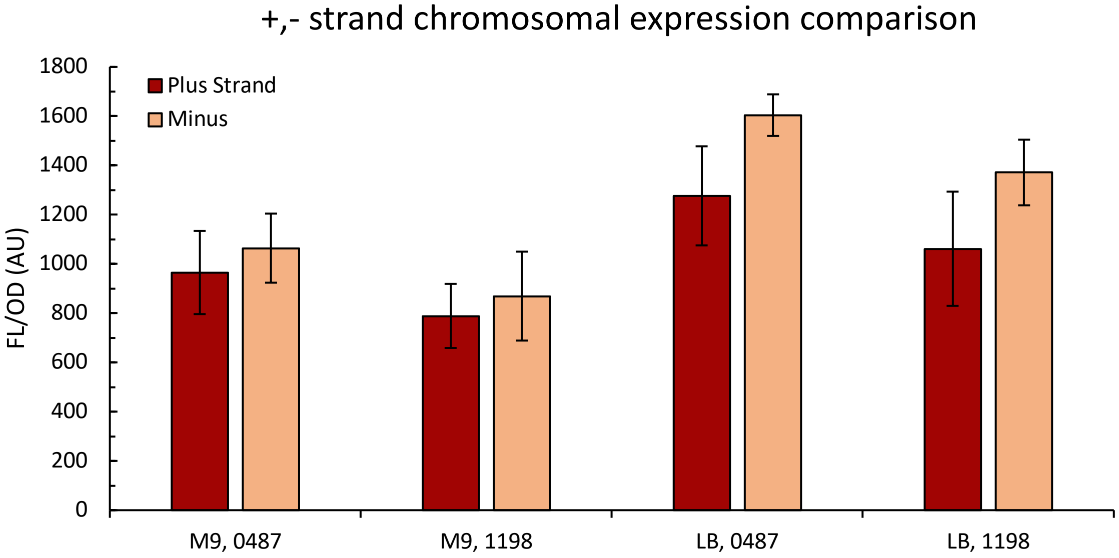

**Supplemental Figure 15.** *Comparison of + vs. - strand expression*. Comparison of + vs. – strand expression at two separate loci in both LB and M9 medium. As can be seen, the – strand generally gives slightly higher expression. 0487 refers to the replacement at ACIAD0487. 1198 refers to the replacement at ACIAD1198. All experiments run in biological triplicate on three separate days (nine replicates total). Error bars are standard deviation.

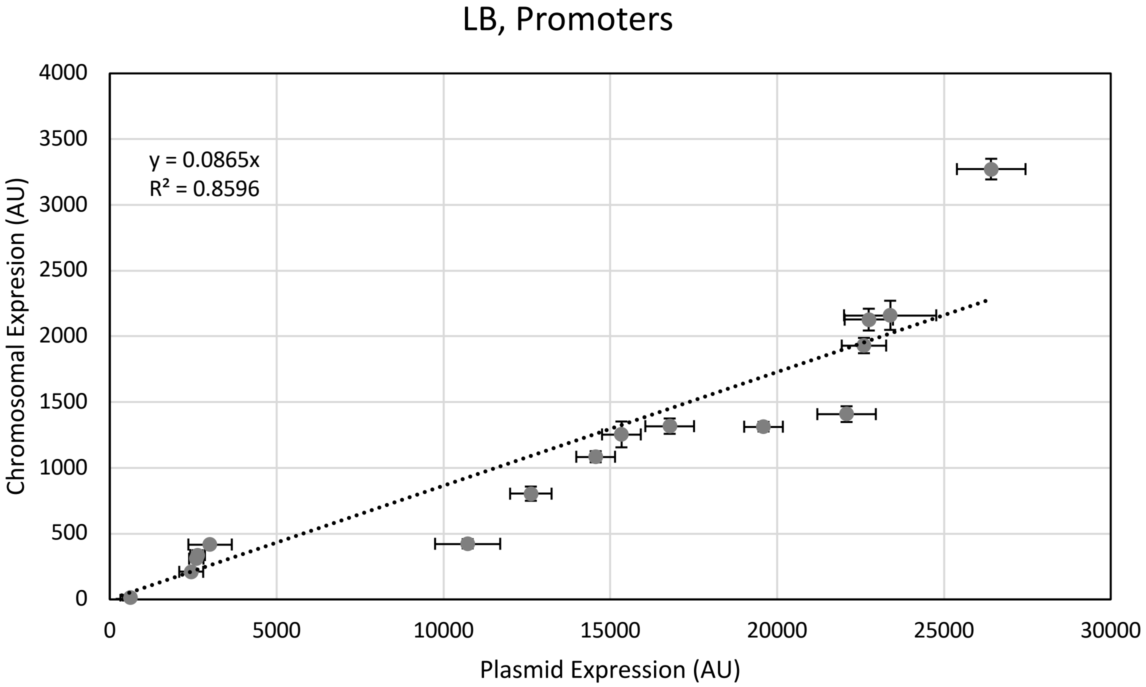

**Supplemental Figure 16.** *Plasmid vs. chromosomally expressed promoter library comparison in LB*. Comparison of the expression for the promoter library on pBAV1k vs. chromosomally integrated. There is fairly obvious trend that in LB the promoter library gives approximately 1/10$^{th}$ the expression at *vanAB* as on the pBAV1k plasmid. All experiments were run in biological triplicate on three separate days (nine replicates total). Error bars are standard error of measurement. Linear regression has intercept set at zero.

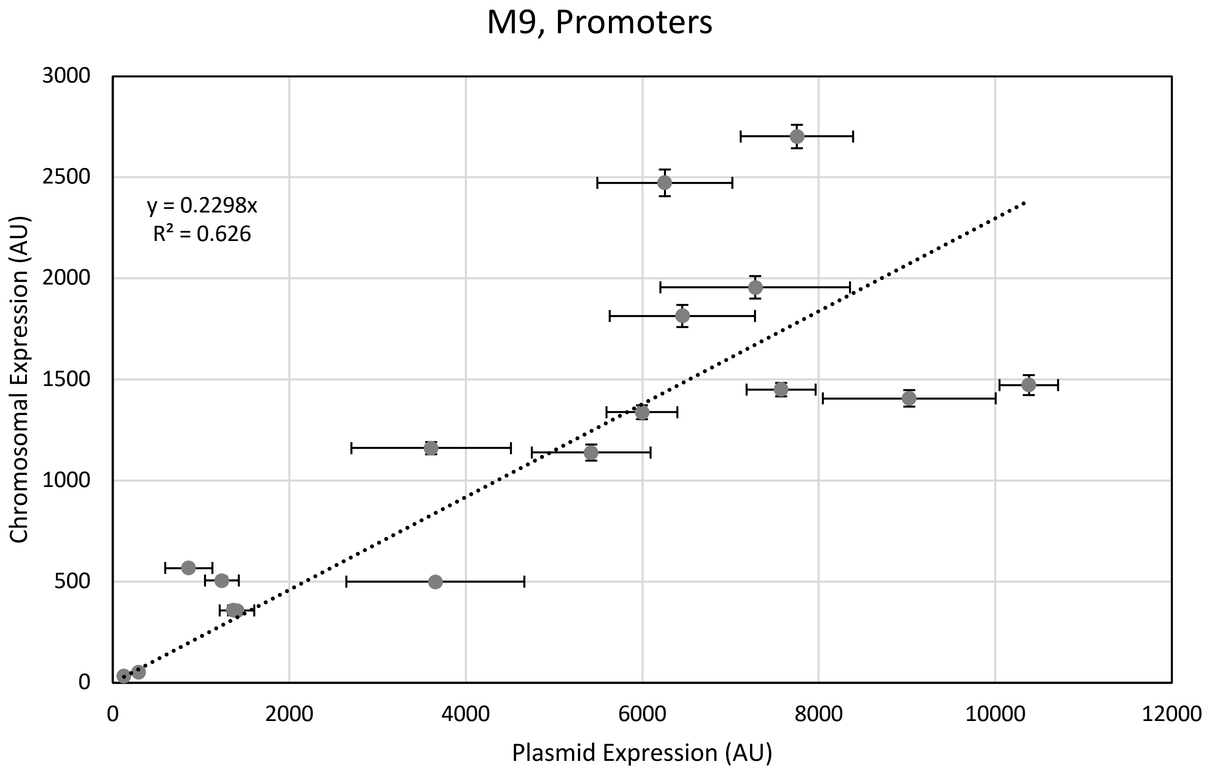

**Supplemental Figure 17.** *Plasmid vs. chromosomally expressed promoter library comparison in M9*. Comparison of the expression for the promoter library on pBAV1k vs. chromosomally integrated. There is modest trend that in M9 the promoter library gives approximately 1/5$^{th}$ the expression at *vanAB* as on the pBAV1k plasmid, but it is not entirely consistent. All experiments were run in biological triplicate on three separate days (nine replicates total). Error bars are standard error of measurement. Linear regression intercept set at zero.

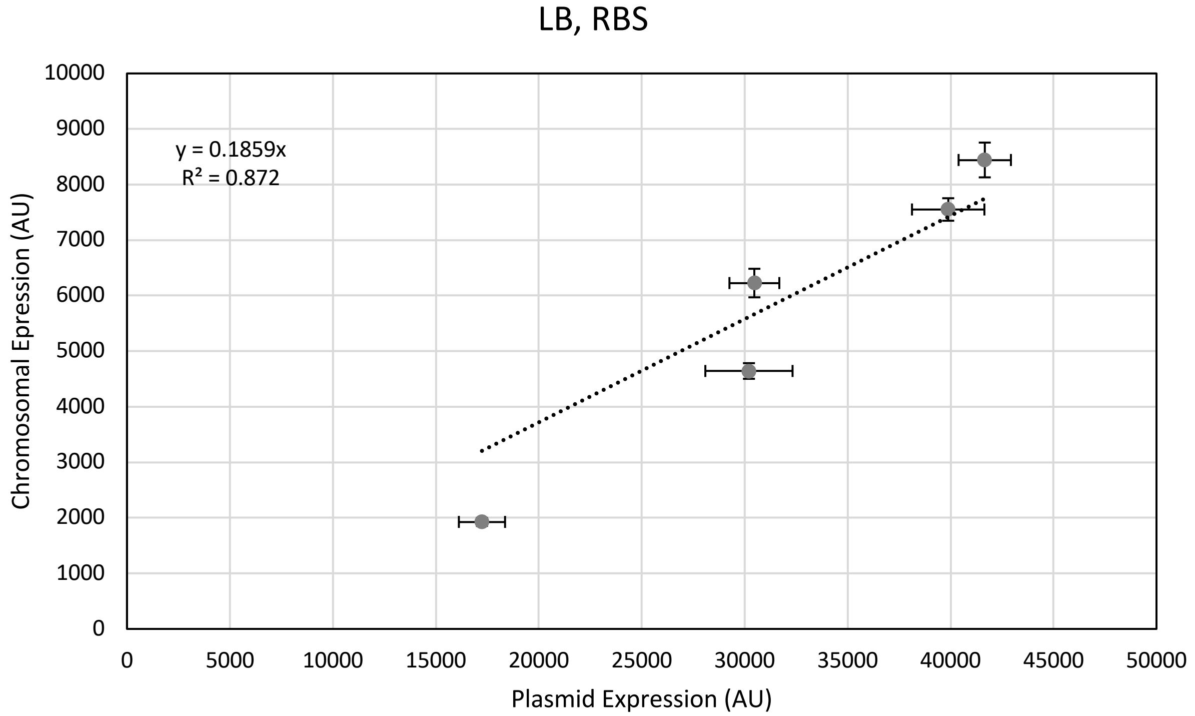

**Supplemental Figure 18.** *Plasmid vs. chromosomally expressed RBS variant comparison in LB.* Comparison of the expression for the RBS/BCD variants on pBAV1k vs. chromosomally integrated. There is trend that in LB the BCD variants gives approximately 1/5$^{th}$ the expression at *vanAB* as on the pBAV1k plasmid. All experiments were run in biological triplicate on three separate days (nine replicates total). Error bars are standard error of measurement. Linear regression intercept set to zero.

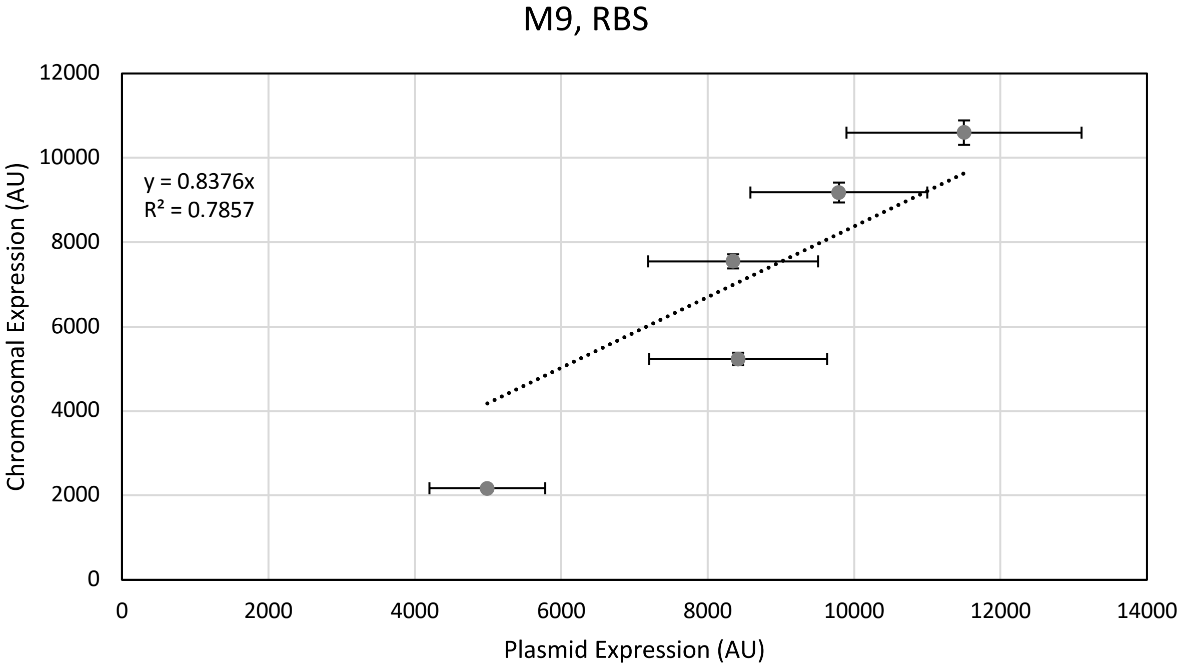

**Supplemental Figure 19.** *Plasmid vs. chromosomally expressed RBS variant comparison in M9*. Comparison of the expression for the RBS/BCD variants on pBAV1k vs. chromosomally integrated. Surprisingly, as can be seen, M9 chromosomal expression of the BCD variants gives over 80% of the expression of the plasmid-based system with modest correlation. All experiments were run in biological triplicate on three separate days (nine replicates total). Error bars are standard error of measurement. Linear regression intercept set at zero.

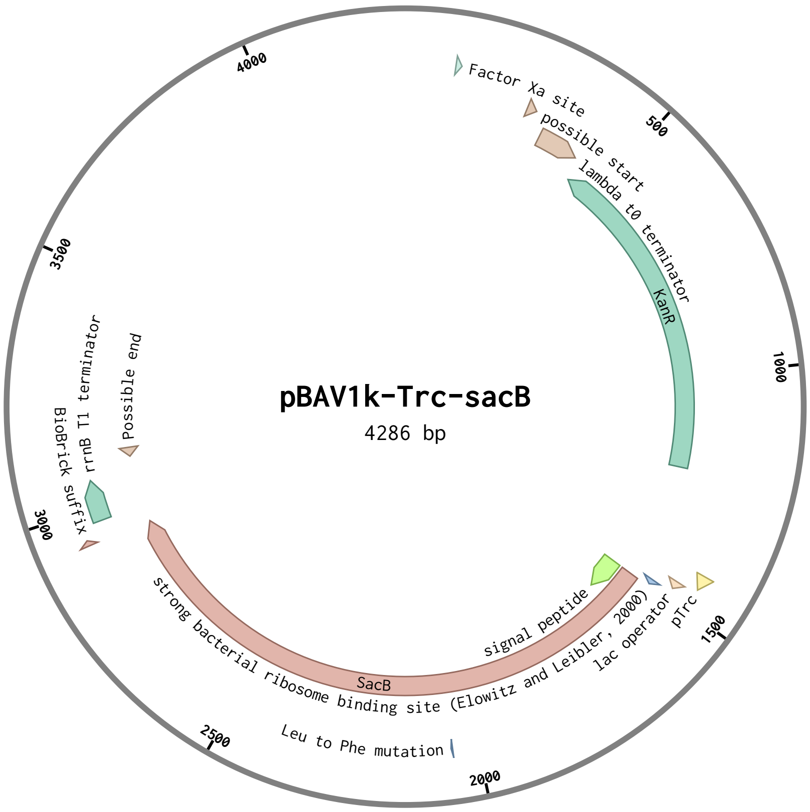

**Supplemental Figure 20.** *Plasmid map for pBWB290 (pBAV1k-Trc-sacB).* (Figure made with Benchling).

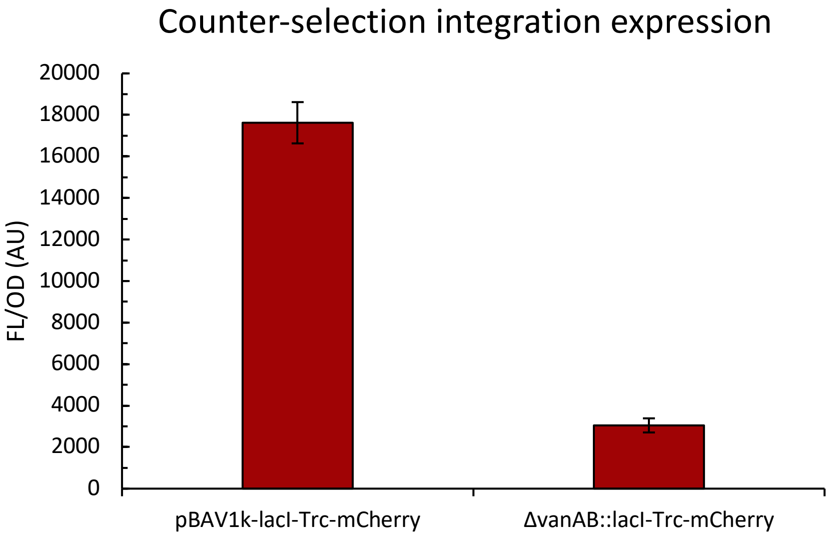

**Supplemental Figure 21.** *Demonstration of chromosomal expression BWB340 (ΔvanAB::lacI-Trc-mCherry).* A marker-less lacI-Trc-mCherry cassette was integrated with counter-selection at ACIAD0980. This vector demonstrates expression from a marker-less cassette. Experiment was run in biological triplicate in 96-well plate in LB. Error bars are standard deviation.

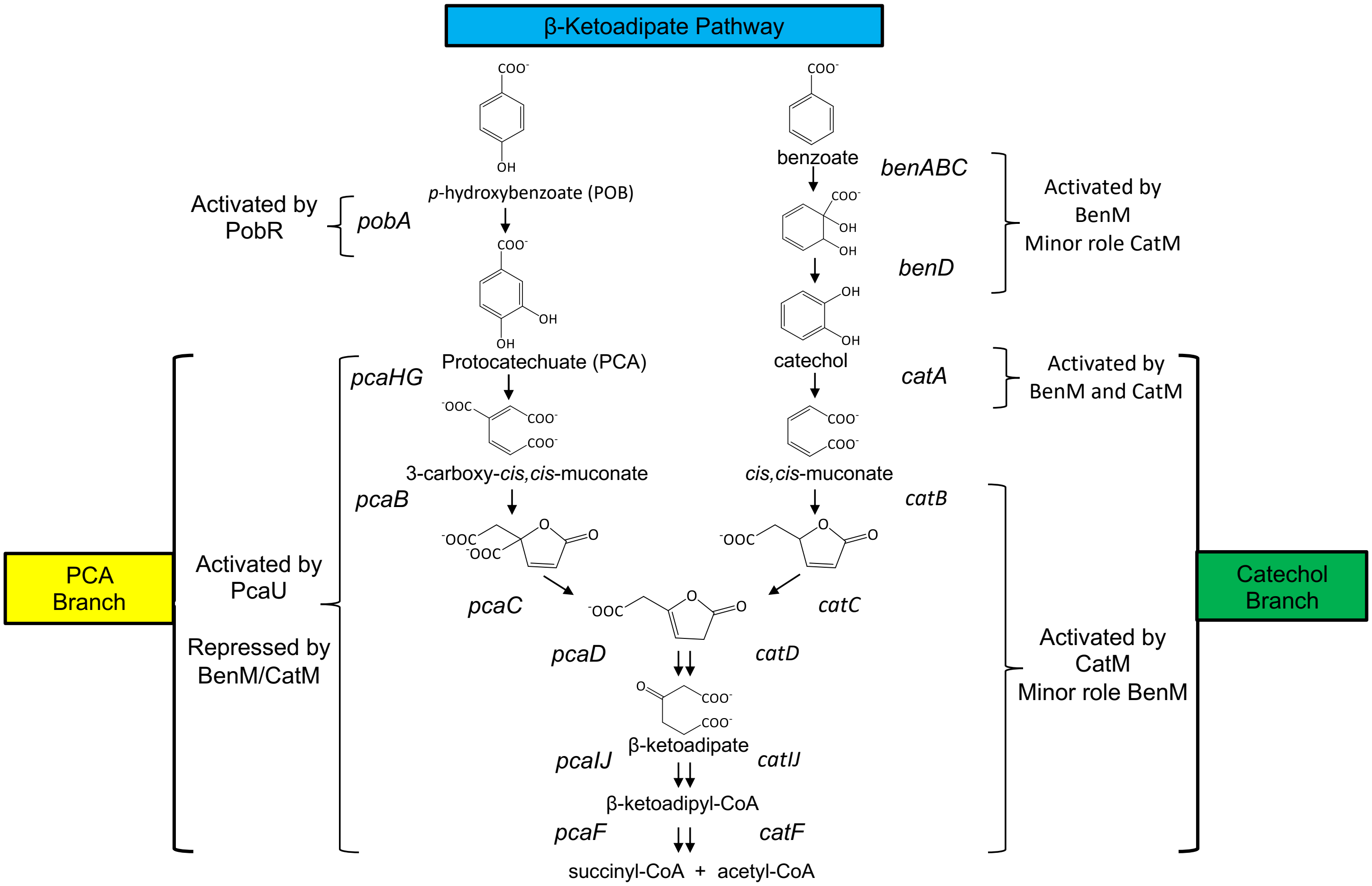

**Supplemental Figure 22**. *The β-ketoadipate pathway of A. baylyi ADP1*. The roles of transcriptional regulators (BenM, CatM, PobR, and PcaU) are indicated. Both BenM and CatM respond to *cis,cis*-muconate. BenM additionally responds to benzoate and can activate transcription synergistically with both effectors. PobR responds to POB. PcaU responds to PCA.

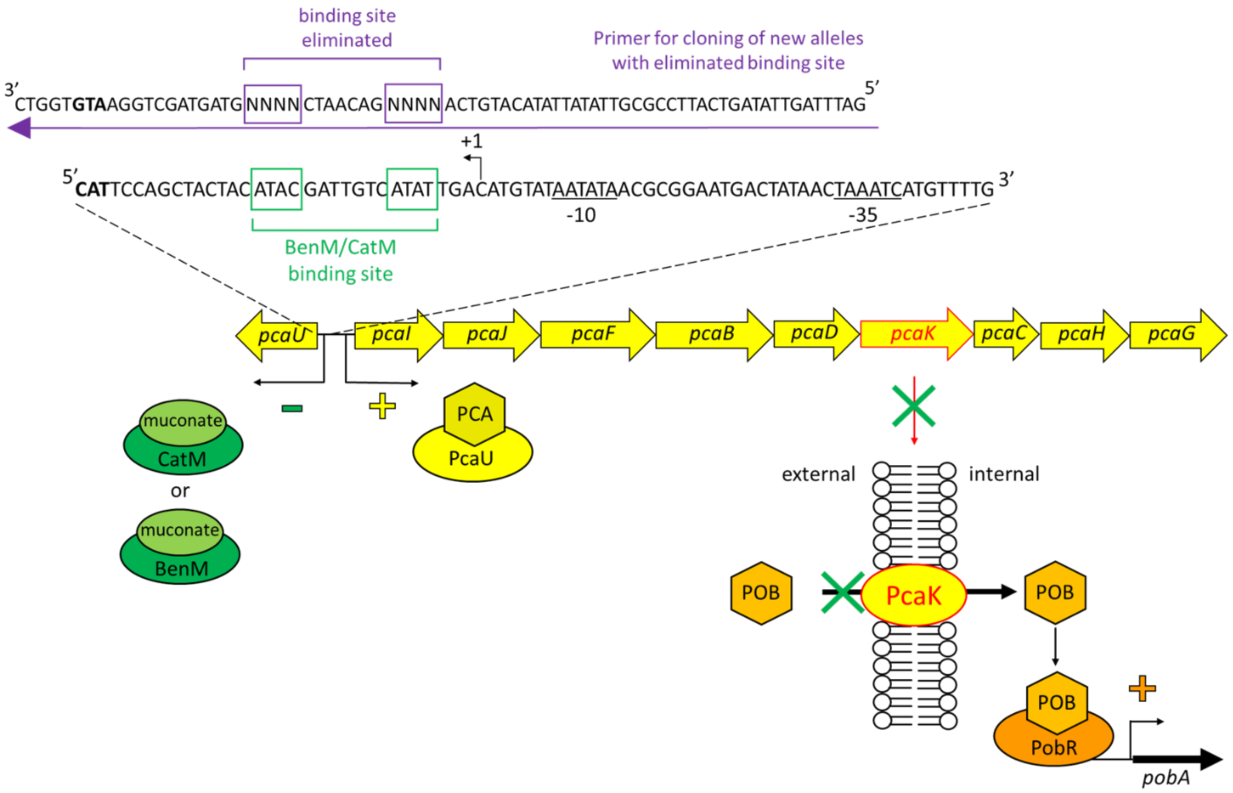

**Supplemental Figure 23***. Regulatory model and experimental approach to prevent BenM and CatM from repressing pca-gene transcription*. Previous studies showed that BenM and CatM bind to the intergenic *pcaU-pcaI* region in a muconate-dependent fashion[10]. Binding of these regulators to the indicated binding site (ATAC-$N_7$-ATAT) could prevent the PcaU activator from being expressed and could reduce transcription of the *pca* operon, which includes *pcaK*. The PcaK transporter can mediate POB uptake to allow the interaction of this effector with the PobR regulator to activate *pobA* transcription. The PobA hydroxylase converts POB to PCA for further catabolism via the PCA branch of the β-ketoadipate pathway. The sequence above the arrow at the top indicates the mutagenic primer used to insert random nucleotides at the 8 important positions in the binding motif.

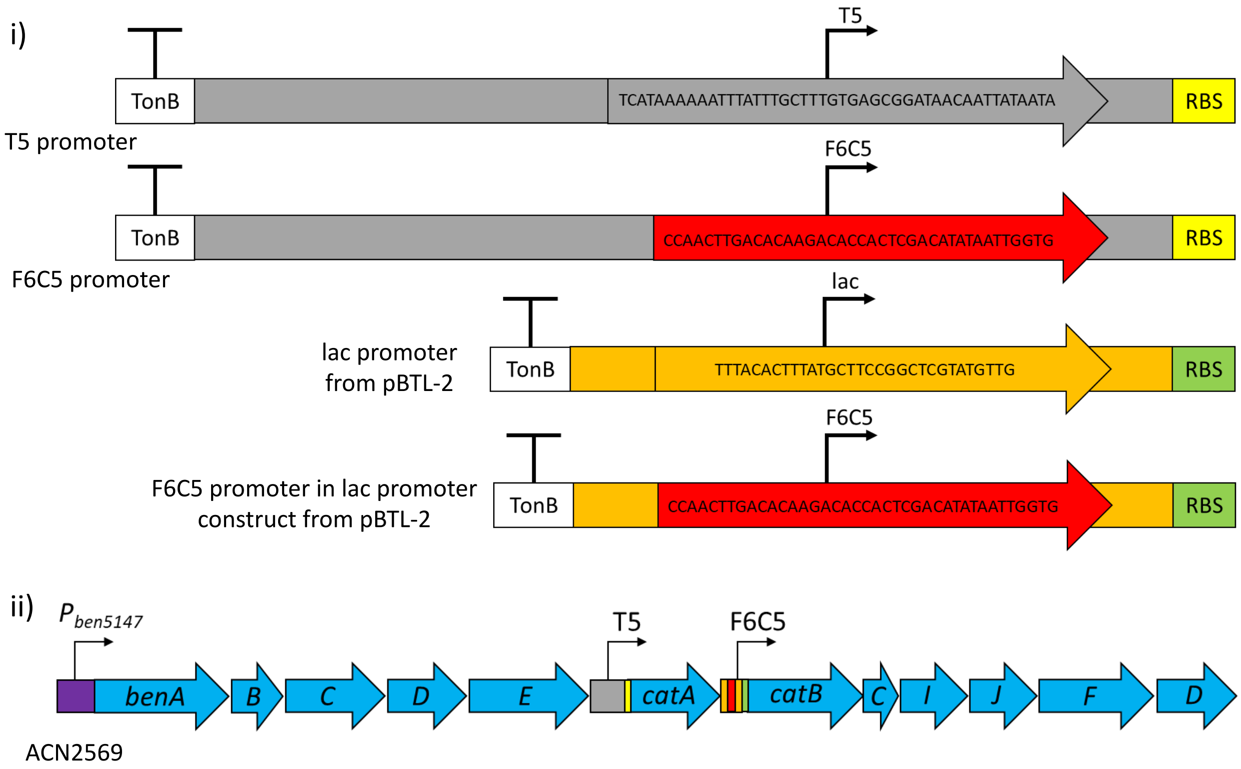

**Supplemental Figure 24.** *Introduction of multiple constitutive promoters in the chromosome to engineer a strain (ACN2569) that grows on benzoate as a carbon source in the absence of the BenM and CatM transcriptional activators*. i) DNA surrounding the key regions of the promoters (arrows) in the synthetic constructs of the tool kit have identical sequences (depicted in gray surrounding the arrows). To avoid homologous recombination between neighboring promoters, the key regions of the F6C5 promoter (red arrow) were used to replace the comparable regions of the *lac* promoter from pBTL-2 so that the DNA surrounding the T5 promoter and that surrounding F6C5 would differ. ii) The T5 promoter was used to transcribe *catA* constitutively and the F6C5 promoter (in the *lac* construct) was used for constitutive transcription of the *catBCIJFD* operon. These expression systems were engineered in the chromosome of a strain that does not encode CatM or BenM and that expresses the *benABCDE* operon from a promoter ($P_{benA5147}$)[14] that does not require an activator.

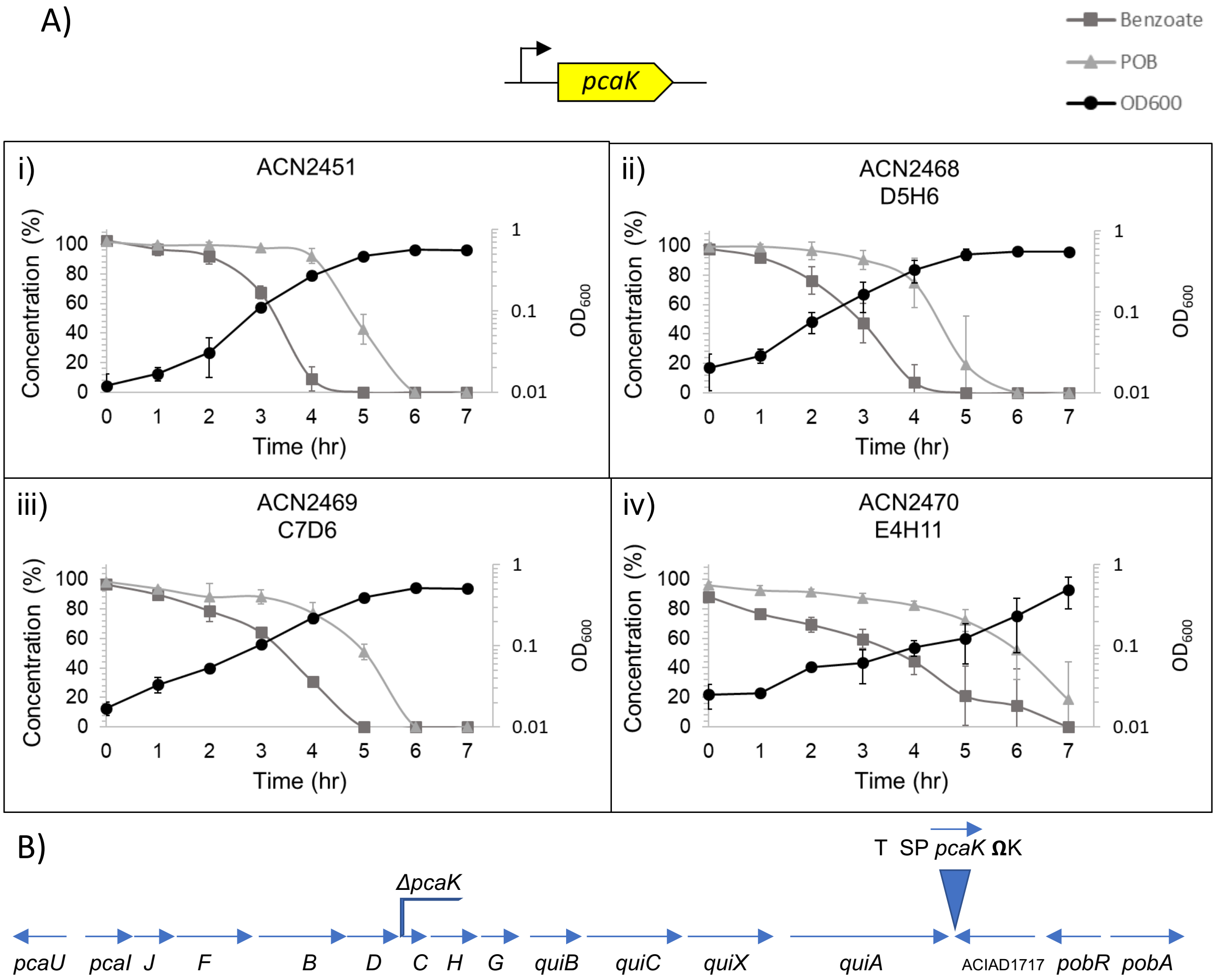

**Supplemental Figure 25.** *Benzoate and POB metabolism by ΔpcaK mutants*. A) Where indicated, *pcaK* was transcribed from a constitutive promoter. Depletion of benzoate and POB (provided as a mixture) was assessed by HPLC of cell-free samples. The 0 time-point is defined as one hour prior to the first detected disappearance of either carbon source. Data points are averages of at least three values with cultures grown on different days. Error bars represent the standard deviation. B) Schematic depiction of the chromosomal regions. In all strains, the *pcaK* coding sequence is deleted from its normal position in an operon. In ACN2468, ACN2469, and ACN2470, *pcaK* was introduced downstream of *quiA* (blue triangle). T represents a terminator insulating *qui*A and *pcaK* transcription. The synthetic promoter (SP) for *pcaK* was either D5H6, C7D6, or E4H11. The ΩK (*52468)* confers kanamycin resistance with the marker flanked by transcriptional and translational stop signals.[15]

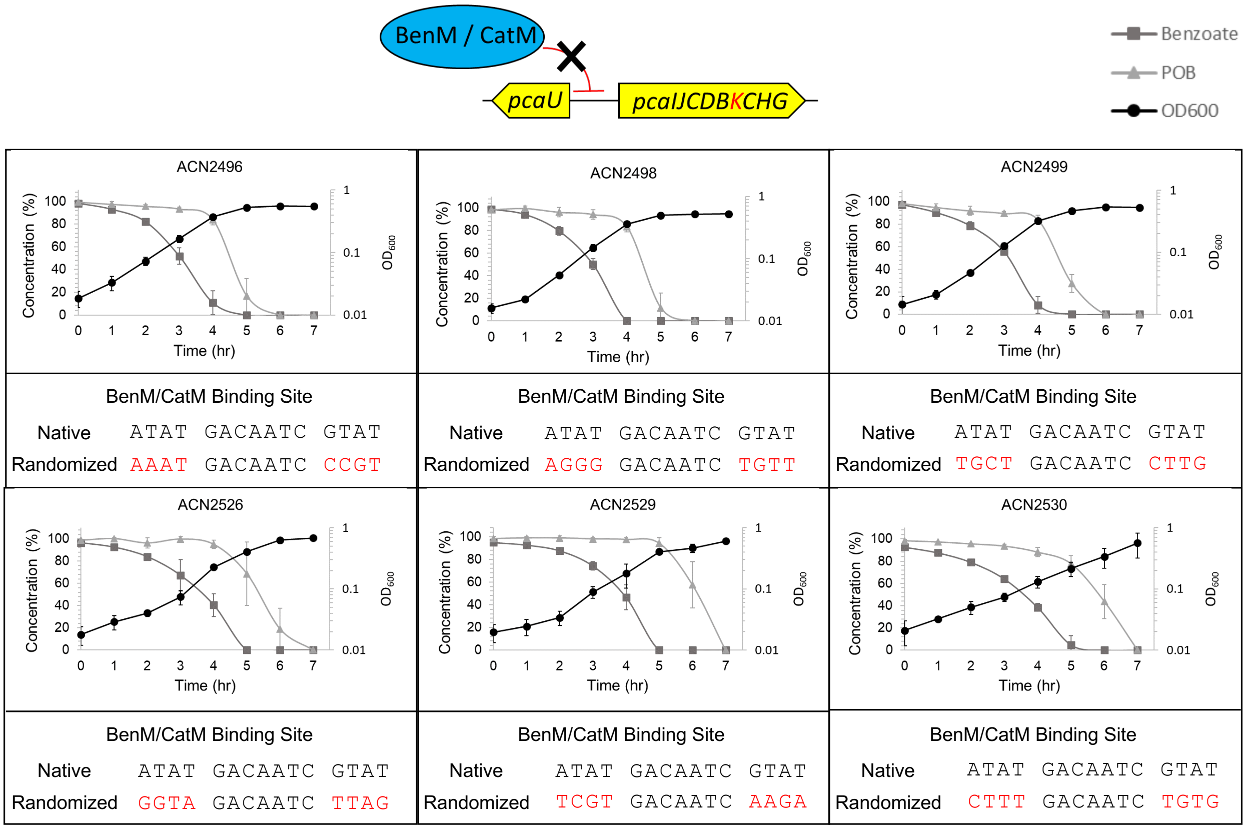

**Supplemental Figure 26.** *Consumption of benzoate and POB by strains with mutations in the pca operator-promoter region*. BenM and CatM bind to the *pcaU-pcaI* intergenic region.[10] To prevent such binding, random changes were made in the consensus sequence used for binding these regulators (ATAT-$N_7$-GTAT). For each strain, red text indicates the mutated sequence in that strain. All strains are able to consume POB and benzoate as sole carbon sources. For data shown here, each strain was provided with a mixture of benzoate and POB in equal amounts. Depletion of these carbon sources, assessed by HPLC analysis of cell-free supernatant fractions of the cultures taken at different times, is shown. The 0 time-point is defined as one hour prior to the first detection of the disappearance of either carbon source. Data points represent the average of at least three values with cultures grown on different days. Error bars represent the standard deviation.

#### Supplemental Tables

| | Conc. (ng/µL) | 260/280 |
|---|---|---|
| WT ADP1 | 7.9 | x |
| pBAV1k, 1 | 199.75 | 1.86 |
| pBAV1k, 2 | 91.03 | 1.93 |
| pBAV1k, 3 | 244.2 | 1.89 |

**Supplemental Table 1.** *ADP1 miniprep yields and purity*. The table shows the yields and purity of using a standard *E. coli*-based miniprep kit and protocol to obtain pBAV1k from ADP1. Minipreps were carried out on 5 mL cultures grown overnight at 30ºC in culture tubes and eluted into 30 µL of nuclease free water. While wild-type ADP1 give no measurable DNA from the miniprep, strains carrying pBAV1k show useful plasmid yield. As noted, ADP1 minipreps require additional spin down steps and time, compared to *E. coli*, after the neutralization step and before binding the plasmid DNA to the column. In general, we ran two separate spins at max speed for 10 minutes, transferring the supernatant to a fresh Eppendorf tube between spins.

| | |
|---|---|
| Trc | AGCTGTTGACAATTAATCATCCGGCTCGTATAATGTGTG |
| C4C9 Trc+ … | TTCACACAGGAAACAGACCAGGGTAAGCTATAATGAGCA |
| F10B12 | CACCGTTGACACTCACGTTCTTGATGTGTATAATGGAAG |
| B2E11 | CCGTGTTGACAATGTGTATCTGTGTTTATATAATGGCCT |
| D7A3 | CCACATTGACAACATCGTATCTGTTCTGTATAATGGTCA |
| C8D2 | CAACTTTGACAGGCACCAACAGTTGGGATATAATCAGGA |
| H1G12 | CATTTTTGACACTAATATTACTTTAATGTATAATGGACG |
| F6C5 | CCAACTTGACACAAGACACCACTCGACATATAATTGGTG |
| B9E2 | GACTTTTGACA--TCGACACACGTGTGGTATAATGGGGT |
| C7D6 | CATCCTTGACACCCCCCCATTTCATCGGTATAATCGGAG |
| C11E8 | CCCACTTGACA-ATTGGCGTACTTTGTATAATATCTC-- |
| B12D11 | CCACATTGACACCCCACTAATGGGGAG-TATAATGTCCC |
| A4F11 | GCTCATTGACACGTTAAATGCTCATTTCTATAATGCGCT |
| D5H6 | GAAATGTGACAGCTTGACATCTGTCATAGTATAATAAGTT |
| E4H11 | TTCCTTTGACACGCACGGGGCGTAATTTTATAATGGTTT |
| D12G6 | CCTGCTTGACAAAGTCCATGTTA-----TAATATGAT– |
| C12H8 | TTCGTGTCGCTCAAGGCGCACTTGGATATATAATAGGCC |

**Supplemental Table 2.** *Table of promoter library sequences*. Table showing the sequences of the promoters in the library, comparing to Trc template. The -35 and -10 boxes are underlined if distinguishable. Promoters are in order of their pBAV1k plasmid-based expression in LB.

| **Supplemental Table 3.** Plasmids for Aromatic Compound Catabolism Studies[a] | | |
|---|---|---|
| pUC18<br>pUC19 | Ap$^R$; cloning vector | [16] |
| pUI1637 | Km$^R$; Source of ΩK cassette | [15] |
| pBTL-2 | Km$^R$; Source of *lac* promoter construct | [9] |
| pBAC1393 | Ap$^R$; ADP1 DNA including *quiA* **(1,721,187-1,723,583)** (MTV580 & MTV581) and ACIAD1717 **(1,723,584-1,724,834)** (MTV582 & MTV583) with engineered SacI recognition site assembled with NEBuilder into pUC18 (MTV584 & MTV585) | This study |
| pBAC1396 | Ap$^R$, Km$^R$ ; ΩK cassette excised from pUI1637 by SacI digestion and inserted into SacI site of pBAC1393 | This study |
| pBAC1561 | Ap$^R$; DNA upstream of the *pca* genes **(1,706,322-1,707,704)** (SRB47 & SRB48), is connected to *pcaK* (**1,713,983-1,715,356)** (SRB51 & SRB52) under the control of the *lac* promoter (from pBTL-2) (SRB49 & SRB50); *pcaK*, is connected to DNA downstream of *pcaG* **(1,717,171-1,718,680)** (SRB53 & SRB54). DNA was assembled with linearized pUC18 using NEBuilder. Note: *pca* regions upstream and downstream of *pcaK* are absent (*pcaUIJFBD* and *pcaCHG,* respectively). | This study |
| pBAC1660 | Ap$^R$; DNA with *trc* promoter controlling mCherry from pBWB162 (SRB121 & SRB122) assembled into pBAC1561 (SRB119 & SRB130) with NEBuilder | This study |
| pBAC1664 | Ap$^R$; DNA with mCherry from pBWB162 (SRB127 & SRB128) assembled into pBAC1561, linearized downstream of *pcaK* (SRB126 & SRB129); A deletion (9$^{th}$ nt of the *pcaK* coding sequence) causes a frameshift and premature truncation | This study |
| pBAC1682 | Ap$^R$; *pcaK*, mCherry, and *aroD* excised from pBAC1664 with EcoRV and DraIII and ligated into the EcoRV and DraIII sites in pBAC1561 | This study |
| pBAC1689 | Ap$^R$; DNA with *pcaDK* **(1,713,160-1,715,372)** (SRB187 & SRB188), *pcaCHG, aroD,* and *quiC* **(1,715,357-1,718,956)** (SRB191 & SRB192), and mCherry (SRB189 & SRB190) assembled in pUC19 (SRB193 & SRB194) with NEBuilder | This study |
| pBAC1690 | Ap$^R$; site directed mutagenesis of pBAC1660 to change *trc* promoter to D5H6 (SRB204 & SRB205) | This study |
| pBAC1702 | Ap$^R$; *pcaK* under the control of the D5H6 promoter replacing *pcaUIJFBDKCHG* **(1,707,705-1,717,170);** PCR fragments containing D5H6 promoter (SRB123 & SRB124), and *pcaK* (SRB125 & SRB126) assembled into pBAC1660 (SRB127 & SRB130) | This study |
| pBAC1717 | Ap$^R$; mCherry removed from pBAC1702 resulting in D5H6-*pcaK52401* (SRB233 & SRB234) | This study |
| pBAC1772 | Ap$^R$, Km$^R$; PCR fragment carrying D5H6-*pcaK52401* from ACN2401 (SRB344 and SRB345) inserted into pBAC1396 (SRB342 and SRB343) | This study |
| pBAC1775 | Ap$^R$; Δ*pcaK52451*<br>*pcaK* removed from pBAC1689 **(1,713,983-1,715,356)** PCR and re-assembled (SRB385 & SRB386) | This study |
| pBAC1779 | Ap$^R$, Km$^R$; P *benE* **(1,438,034-1,439,233)** (SRB330 & SRB341) and *catBCIJFD* **(1,444,700-1,449,606)** (SRB337 & SRB340), and ΩK (SRB338 & SRB339) assembled in pUC19 with (SRB328 & SRB329) | This study |

| Supplemental Table 3 (continued). Plasmids for Aromatic Compound Catabolism Studies[a] | | |
|---|---|---|
| pBAC1780 | Ap$^R$, Km$^R$; site directed mutagenesis of pBAC1772 to change D5H6 promoter to C7D6 promoter controlling *pcaK52469* (SRB389 & SRB390) | This study |
| pBAC1791 | Ap$^R$, Km$^R$; site directed mutagenesis of pBAC1772 to change D5H6 promoter to E4H11 promoter controlling *pcaK52470* (SRB387 & SRB388) | This study |
| pBAC1793 | Ap$^R$, Km$^R$; T5 promoter from pBWB162 (SRB332 & SRB333), *catA* **(1,439,848-1,440,783)** (SRB334 & SRB335) assembled in pBAC1779 (SRB331 & SRB336) to make T5-*catA-catBCIJFD52528* | This study |
| pBAC1794 | Ap$^R$, Km$^R$; *pcaU52529;* **1,709,012-1,707,109** (*pcaU* coding sequence with adjacent mutated BenM/CatM binding site[b]:**,** TCGTGACAATCAAGA**)** (SRB368 & SRB369), with ΩK (SRB375 & SRB376) downstream of *pcaU* (SRB377 & SRB378), and with *pcaI* region in native position **(1,708,973-1,709,914)** (SRB370 & SRB371) assembled in pUC19 (SRB372 & SRB373) | This study |
| pBAC1795 | Ap$^R$, Km$^R$; *pcaU52496;* **1,709,012-1,707,109** (*pcaU* coding sequence with adjacent mutated BenM/CatM binding site[b]: AAATGACAATCCCGT) (SRB368 & SRB369), with ΩK (SRB375 & SRB376) downstream of *pcaU* (SRB377 & SRB378), and with *pcaI* region in native position **(1,708,973-1,709,914)** (SRB370 & SRB371) assembled in pUC19 (SRB372 & SRB373) | This study |
| pBAC1797 | Ap$^R$, Km$^R$; *pcaU52498;* **1,709,012-1,707,109** (*pcaU* coding sequence with adjacent mutated BenM/CatM binding site[b]: AGGGGACAATCTGTT) (SRB368 & SRB369), with ΩK (SRB375 & SRB376) downstream of *pcaU* (SRB377 & SRB378), and with *pcaI* region in native position **(1,708,973-1,709,914)** (SRB370 & SRB371) assembled in pUC19 (SRB372 & SRB373) | This study |
| pBAC1798 | Ap$^R$, Km$^R$; *pcaU52530* **1,709,012-1,707,109** (*pcaU* coding sequence with adjacent mutated BenM/CatM binding site[b]: CTTTGACAATCTGTG) (SRB368 & SRB369), with ΩK (SRB375 & SRB376) downstream of *pcaU* (SRB377 & SRB378), and with *pcaI* region in native position **(1,708,973-1,709,914)** (SRB370 & SRB371) assembled in pUC19 (SRB372 & SRB373) | This study |
| pBAC1799 | Ap$^R$, Km$^R$; *pcaU52526* **1,709,012-1,707,109** (*pcaU* coding sequence with adjacent mutated BenM/CatM binding site[b]: GGTAGACAATCTTAG) (SRB368 & SRB369), with ΩK (SRB375 & SRB376) downstream of *pcaU* (SRB377 & SRB378), and with *pcaI* region in native position **(1,708,973-1,709,914)** (SRB370 & SRB371) assembled in pUC19 (SRB372 & SRB373) | This study |
| pBAC1800 | Ap$^R$, Km$^R$; *pcaU52499* **1,709,012-1,707,109** (*pcaU* coding sequence with adjacent mutated BenM/CatM binding site[b]: TGCTGACAATCCTTG) (SRB368 & SRB369), with ΩK (SRB375 & SRB376) downstream of *pcaU* (SRB377 & SRB378), and with *pcaI* region in native position **(1,708,973-1,709,914)** (SRB370 & SRB371) assembled in pUC19 (SRB372 & SRB373) | This study |
| pBAC1805 | Ap$^R$, Km$^R$; Modification of pBAC1793 to add ADP1 DNA downstream of ΩK; additional DNA **(1,449,607-1,450,626)** (SRB404 & SRB405) was integrated into pBAC1793 (SRB406 & SRB407) including T5-*catA-catBCIJFD52528* | This study |
| pBAC1817 | Ap$^R$, site directed mutagenesis of pBAC1682 to change *lac* promoter to F6C5 promoter controlling *pcaK* (SRB423 & SRB424) | This study |

| **Supplemental Table 3 (continued).** Plasmids for Aromatic Compound Catabolism Studies[a] | | |
|---|---|---|
| pBAC1829 | Ap$^R$, Km$^R$; The F6C5 replaces the *lac* promoter from pBTL-2 (SRB441 & SRB442) assembled upstream of *catBCIJFD* in pBAC1805 (SRB439 & SRB440) | This study |

a. Bold numbers correspond to positions on the ADP1 chromosome in NCBI entry NC_005966; PCR primers used to generate DNA fragments are underlined and in parentheses; ΩK refers to the omega cassette from pUI1637 encoding Km$^R$; Unless otherwise stated, plasmids were assembled in *E. coli* XLI Blue by the method described by Kostylev et al.[5]

b. When the BenM/CatM binding site was mutated, the specific sequence on the plasmid is indicated, and nucleotides in red text mark mutations that alter the wild-type sequence, which is ATATGACAATCGTAT (**1,708,958- 1,708,972).**

| **Supplemental Table 4.** ADP1-Derived Strains for Aromatic Compound Catabolism Studies[a] | | |
|---|---|---|
| ADP1 | Wild type (BD413) | 1, 2 |
| ACN825 | *benM*::Ω*S4036*, *benA5147*, *Δcat5825*(*catA-catD*); A promoter mutation in *benA* ($P_{benA5147}$) changes G to A at **1,433,999** causing high-level transcription without requiring activation by BenM or CatM | 7 |
| ACN2067 | Δ*pcaCHG52067;* Deletion from **1,715,357- 1,718,112.** Donor DNA replaced the wild-type genomic region with a deletion via homologous recombination events in two locations, one in *pcaK* and the other downstream of *pcaG.* pBAC1561/AatII X ADP1; screened for the loss of ability to grow on POB | This study |
| ACN2451 | Δ*pcaK52451;* Deletion of *pcaK* (**1,713,983 – 1, 715,356**), also restored wild-type sequence in the recipient strain, *pcaCHG* (**1,715,357 – 1,718,112);** pBAC1775/AatII X ACN2067; selected by growth on POB | This study |
| ACN2468 | D5H6-*pcaK52468*; Ω*K52468*; Δ*pcaK52451;* ΩK downstream of ACIAD1717 *pcaK* under the control of the D5H6 promoter after **1,723,583** and deleted from its native location; pBAC1772/AatII X ACN2451; selected by Km$^R$ | This study |
| ACN2469 | C7D6-*pcaK52469*; Ω*K52468*; Δ*pcaK52451;* ΩK downstream of ACIAD1717 *pcaK* under the control of the C7D6 promoter after **1,723,583** and deleted from its native location; pBAC1780/AatII X ACN2451; selected by Km$^R$ | This study |
| ACN2470 | E4H11-*pcaK52470*; Ω*K52468*; Δ*pcaK52451;* ΩK downstream of ACIAD1717 *pcaK* under the control of the E4H11 promoter after **1,723,583** and deleted from its native location; pBAC1791/AatII X ACN2451; selected by Km$^R$ | This study |
| ACN2471 | H1G12-*pcaK52471*; Ω*K52468*; Δ*pcaK52451;* ΩK downstream of ACIAD1717 *pcaK* under the control of the H1G12 promoter after **1,723,583** and deleted from its native location; Strain unable to grow on benzoate or POB for unknown reason, but likely related to high PcaK expression pBAC1792/AatII X ACN2451; selected by Km$^R$ | This study |
| ACN2496 | *pcaU52496*; Ω*K52496* (*pcaU* coding sequence with adjacent mutated BenM/CatM binding site[b]: AAATGACAATCCCGT); ΩK downstream of *pcaU* pBAC1795/AatII X ADP1; selected by Km$^R$ | This study |

| Supplemental Table 4 (continued). ADP1-Derived Strains for Aromatic Compound Catabolism Studies[a] | | |
|---|---|---|
| ACN2498 | *pcaU52498*; ΩK*52496* (*pcaU* coding sequence with adjacent mutated BenM/CatM binding site[b]: AGGGGACAATCTGTT); ΩK downstream of *pcaU* pBAC1797/AatII X ADP1; selected by Km[R] | This study |
| ACN2499 | *pcaU52499* ΩK*52496* (*pcaU* coding sequence with adjacent mutated BenM/CatM binding site[b]: TGCTGACAATCCTTG); ΩK downstream of *pcaU* pBAC1800/AatII X ADP1; selected by Km[R] | This study |
| ACN2526 | *pcaU52526;* ΩK*52496* (*pcaU* coding sequence with adjacent mutated BenM/CatM binding site[b]: GGTAGACAATCTTAG); ΩK downstream of *pcaU* pBAC1799/AatII X ADP1; selected by Km[R] | This study |
| ACN2529 | *pcaU52529;* ΩK*52496* (*pcaU* coding sequence with adjacent mutated BenM/CatM binding site[b]: TCGTGACAATCAAGA**);** ΩK downstream of *pcaU* pBAC1794/AatII X ADP1; selected by Km[R] | This study |
| ACN2530 | *pcaU52530;* ΩK*52496* (*pcaU* coding sequence with adjacent mutated BenM/CatM binding site[b]: CTTTGACAATCTGTG); ΩK downstream of *pcaU* pBAC1798/AatII X ADP1; selected by Km[R] | This study |
| ACN2569 | T5-*catA52569; F6C5-catBCIJFD52569*; ΩK*52569*; *benM*::ΩS*4036*, *benA5147;* Δ*catM5293; catA* **(1,439,848-1,440,783)** controlled by the T5 promoter and *catBCIJFD* **(1,444,715-1,449,246)** controlled by the F6C5 promoter; pBAC1829 X ACN825[c]; selected by Km[R] | This study |

a. *A. baylyi* strains were derived from ADP1, previously known as *Acinetobacter calcoaceticus* or *Acinetobacter* sp.[2]; bold numbers correspond to positions on the ADP1 chromosome in NCBI entry NC_005966.; ΩK and ΩS indicate cassettes that confer resistance to kanamycin or streptomycin and spectinomycin;[15] Underlined text indicates the donor DNA and, where relevant, the restriction enzyme used to linearize a plasmid (pBAC number/Enzyme). The donor DNA was used to transform (X) the indicated recipient strain.

b. When the BenM/CatM binding site was mutated, the specific sequence on the chromosome is indicated, and nucleotides in red text mark mutations that alter the wild-type sequence, which is ATATGACAATCGTAT (**1,708,958- 1,708,972).**

### Supplemental Note

##### Aromatic Carbon Source Preferences in ADP1

While the molecular basis for hierarchical consumption remains unclear, in a two-compound mixture, POB degradation commences only after the majority of benzoate is consumed. This regulation is so inflexible that if benzoate and POB are provided to certain mutants that cannot use benzoate as the carbon source, POB consumption is completely prevented[10, 14, 17]. These mutants, with blocks in the catechol branch, fail to grow when provided with POB and benzoate together, even though POB alone permits growth and there are no mutations in genes encoding any of the necessary enzymes for POB catabolism. However, POB consumption is only inhibited if benzoate can be endogenously metabolized to muconate, consistent with this metabolite serving as a signaling molecule[17]. Two muconate-responsive transcriptional regulators, BenM and CatM, have been implicated in cross regulation of both branches of the β-ketoadipate pathway in ADP1[10]. In response to muconate, they activate the transcription of *ben* and *cat* genes, needed for benzoate and catechol catabolism. In contrast, they appear to repress PCA degradation, as evidenced by muconate-dependent binding to DNA between the divergent *pcaU* and *pcaI* genes[10]. The location of the binding site (**Supplemental Fig. 23**) suggests that PCA catabolism is prevented due to the repressed expression of a transcriptional activator (PcaU) and/or by affecting the operator-promoter region of the *pcaIJFBDKCHG* operon. PcaU is needed to activate the transcription of this operon. Binding of BenM and CatM would not directly explain how the conversion of POB to PCA, mediated by PobA, is prevented. During growth in the presence of benzoate, the transcription of *pobA* is inhibited[10], but there is no proximal site near this gene for binding BenM and CatM to account for the effect. While repression by BenM and CatM may lower the uptake of POB by PcaK, another possibility is that repressed expression of PcaU affects the ability of this regulator to control the transcription of additional genes in a larger, but as yet unknown, PcaU regulon.

##### Effects of Transcribing *pcaK* from Synthetic Promoters on Co-Consumption of Benzoate and POB

The known inhibition of *pobA* transcription could result from lowered POB uptake in the presence of benzoate, because *pobA* transcription depends on POB to interact with the PobR regulator (**Supplemental Fig. 23**). This type of inducer exclusion, which is a canonical mechanism of catabolite repression, might be overcome by increasing transcription of *pcaK*. To test this possibility, *pcaK* was deleted from its native position in the *pca* operon, where it can be repressed by BenM and CatM. Transcription of *pcaK* elsewhere in the chromosome was engineered under the constitutive control of three synthetic promoters of different strengths (D5H6 in ACN2468, C7D6 in ACN2469, and E4H11 in ACN2470). However, simultaneous consumption of benzoate and POB does not result when *pcaK* is transcribed from any of these promoters. The consumption patterns all resemble that of ACN2451, where the *pcaK* gene is absent **(Fig. 5bii** and **Supplemental Fig. 25**). These results suggest that low PcaK expression is insufficient to explain the preferential consumption of benzoate. However, inducer exclusion cannot be ruled out as a mechanism of regulation. Additional transporters, such as VanK, can also import POB, and little is known about the regulation, interplay, and relative importance of these proteins for

POB uptake in the presence of benzoate[18, 19]. Moreover, further study is needed to confirm that the change in transcriptional control of *pcaK* actually results in increased uptake of POB. While the preference for benzoate did not change, adverse effects on growth may occur with increasing transcription of *pcaK*. The generation times for the two strains with the strongest promoters (ACN2469 and ACN2470) were approximately 60 minutes, a value slightly higher than those of the others (ADP1, ACN2451, and ACN2468), which were all close to 50 minutes. Strain ACN2470 had a significantly longer lag period than the other strains, approximately 8 hours compared to 3-4 hours for the others. Furthermore, a comparable strain (ACN2471) constructed with an even stronger promoter, H1G12, failed to grow on benzoate or POB. These results highlight the importance of having a range of promoter strengths available for different experiments.

**Effects of Removing the BenM-CatM Binding Site on Co-Consumption of Benzoate and POB**

A second approach was used to determine whether the binding of BenM or CatM to DNA in the *pca*-region is sufficient to account for the preferential consumption of benzoate. Mutagenesis was used to eliminate sequences used to bind BenM and CatM while leaving intact the DNA needed for transcription of the divergently oriented *pcaU* and *pcaI* genes. The consensus binding site for BenM and CatM is ATAC-$N_7$-GTAT, and a sequence with one mismatch (ATAC-$N_7$-ATAT, **Supplemental Fig. 23**) is likely to bind a regulatory protein dimer in the *pca* region. The most important sequence features are T-$N_{11}$-A surrounded by a small region of dyad symmetry. To remove these key features while minimizing adverse effects on *pca* transcription, only the ATAC and ATAT sequences were randomized using primers synthesized to contain any nucleotide at these positions (see Methods). After DNA with random substitutions in these 8 nucleotide positions had been integrated in the chromosome by allelic replacement, strains that can grow on POB as the sole carbon source were selected. Several strains were further characterized and tested for the ability to consume mixtures of benzoate and POB. All of these strains (ACN2496, ACN2498, ACN2499, ACN2526, ACN2529, and ACN2530) had lost the important features needed to bind BenM and CatM, yet all consumed benzoate before POB (**Supplemental Fig. 26**). Consistent with the studies above using synthetic promoters, these efforts to eliminate BenM and CatM binding to *pca* DNA indicate that repression of *pca*-operon expression (including *pcaK*) is insufficient to explain the preferential consumption of benzoate. Furthermore, the results with these binding site mutations suggest that repression of PcaU expression is insufficient to explain the benzoate and POB consumption pattern.

**Effects of Removing BenM and CatM on Co-Consumption of Benzoate and POB**

Strains lacking BenM and CatM were previously shown to consume benzoate and POB simultaneously[10]. We sought to conduct similar experiments in a different fashion, as explained below. Growth on benzoate as the carbon source requires high-level transcription of the *benABCDE* operon, the *catA* gene, and the *catBCIJFD* operon. In previous studies, in strains lacking BenM and CatM, *ben*-gene transcription was increased with a spontaneous promoter mutation that obviates the need for the native transcriptional activators. To increase transcription of the *cat* genes, large sections of the corresponding chromosomal regions were amplified in a tandem array so that many copies of weak promoters would result in high-level *cat*-gene transcription. For the current project, we generated a differently constructed strain to grow on benzoate without BenM and CatM for two reasons. First, the amplification of the *cat* genes, which occurred in previous studies, requires substantial changes to the chromosome that might skew experimental results. Therefore, to confirm the effects of BenM and CatM-independent growth patterns a new strain was constructed with a single chromosomal copy of the *ben* and *cat* genes. Second, for biotechnology applications, such as lignin valorization, strains with regional gene amplification are undesirable because of genetic instability. The new genetic tool kit enables high-level transcription of *catA* and the *catBCIJFD* operon using synthetic promoters. Strain ACN2569, which does not encode either BenM or CatM, was engineered to transcribe *catA* from the T5 promoter and to transcribe the *catBCIJFD* operon from the F6C5 promoter (**Supplemental Fig. 24**). These synthetic promoters together with $P_{benA!5147}$, which is the same promoter mutation that was used previously for high *ben*-operon expression, enable ACN2569 to growth on benzoate as the sole carbon source. Consistent with previous results using strains with chromosomal gene amplification, the absence of BenM and CatM allow benzoate and POB to be co-consumed (**Figure 5**).

**Concluding Remarks**

Understanding cross regulation of the two branches of the β-ketoadipate pathway in ADP1 has important implications for the co-metabolism of aromatic compounds in mixed feedstocks. This topic is important for fundamental understanding of regulation, metabolic engineering, and applications such as lignin valorization and plastics recycling. The new genetic tools described in this report are enabling long-discussed regulatory hypotheses to be experimentally tested. Our data indicate that while BenM and CatM play an important role in the preferential consumption of benzoate before POB, this regulation cannot be explained solely by an ability to bind upstream of the *pcaU* gene. BenM and CatM participate in a complex regulon and have varied, and overlapping, roles in both transcriptional activation and repression. Future studies are needed to characterize this regulon fully. Nevertheless, strain ACN2569 consumes benzoate and POB at the same time, and it can serve as a starting strain for further improvements to optimize the consumption of lignin-derived mixtures.

#### Supplemental Files Description

Accompanying this work are three Excel files, along with sequence files for the tools plasmids and chromosomal integrations. **Supplemental File 1** contains the strains created in the tools portion of the work, with strain numbers and descriptions. **Supplemental File 2** contains all the primers created in this study. **Supplemental File 3** contains all the plasmids created for the tools portion of the work, with plasmid numbers and descriptions. In addition, there are GenBank files for the tools constructs in a folder structure.
